# Identifying cell states in single-cell RNA-seq data at statistically maximal resolution

**DOI:** 10.1101/2023.10.31.564980

**Authors:** Pascal Grobecker, Erik van Nimwegen

## Abstract

Single-cell RNA sequencing (scRNA-seq) has become a popular experimental method to study variation of gene expression within a population of cells. However, obtaining an accurate picture of the diversity of distinct gene expression states that are present in a given dataset is highly challenging because the sparsity of the scRNA-seq data and its inhomogeneous measurement noise properties. Although a vast number of different methods is applied in the literature for clustering cells into subsets with ‘similar’ expression profiles, these methods generally lack rigorously specified objectives, involve multiple complex layers of normalization, filtering, feature selection, dimensionality-reduction, employ *ad hoc* measures of distance or similarity between cells, often ignore the known measurement noise properties of scRNA-seq measurements, and include a large number of tunable parameters. Consequently, it is virtually impossible to assign concrete biophysical meaning to the clusterings that result from these methods.

Here we address the following problem: Given raw unique molecule identifier (UMI) counts of an scRNA-seq dataset, partition the cells into subsets such that the gene expression states of the cells in each subset are statistically indistinguishable, and each subset corresponds to a distinct gene expression state. That is, we aim to partition cells so as to maximally reduce the complexity of the dataset without removing any of its meaningful structure. We show that, given the known measurement noise structure of scRNA-seq data, this problem is mathematically well-defined and derive its unique solution from first principles. We have implemented this solution in a tool called Cellstates which operates directly on the raw data and automatically determines the optimal partition and cluster number, with zero tunable parameters.

We show that, on synthetic datasets, Cellstates almost perfectly recovers optimal partitions. On real data, Cellstates robustly identifies subtle substructure within groups of cells that are traditionally annotated as a common cell type. Moreover, we show that the diversity of gene expression states that Cellstates identifies systematically depends on the tissue of origin and not on technical features of the experiments such as the total number of cells and total UMI count per cell. In addition to the Cellstates tool we also provide a small toolbox of software to place the identified cellstates into a hierarchical tree of higher-order clusters, to identify the most important marker genes at each branch of this hierarchy, and to visualize these results.

## 1 Introduction

All cells in multicellular organisms contain the same genome with typically around 20,000 genes but are able to take on a wide variety of phenotypes and perform specialized functions by selective expression of these genes. Therefore, it is one of the fundamental problems of cell biology to characterize the gene expression states cells take on in a multicellular organism. Addressing this requires investigating gene expression states in single cells, which has become possible through progress in the development of single-cell technologies, and single-cell RNA sequencing (scRNA-seq) in particular, over the last years. Numerous cell atlas projects [1, 2, 3, 4, 5] using this approach are already published or in progress. It is often assumed that cells can be divided into discrete cell types which have characteristic molecular profiles and perform specific functions, but despite of all the available experimental data, it is still debated how such discrete types should be defined [6, 7, 8, 9]. Indeed, it is also often proposed that gene expression states are not discrete but rather occupy a continuous subspace of gene expression space, which is typically assumed to be of much lower dimensionality than the full gene expression space [10, 11].

It is thus currently not clear to what extent the assumption that cells can be grouped into discrete states is appropriate. Arguably, during cellular differentiation cells must be traversing an approximately continuous space of gene expression states, but for fully differentiated tissues it may not be unreasonable to approximate cells as deriving from a set of discrete states. However, even if we take for granted the assumption that cells take on discrete states or ‘types’, there is currently also no agreement regarding how such types should be defined or identified. That is, although intuitively cells of the same type should have ‘similar’ expression profiles, there is currently no agreed upon metric of closeness of gene expression states and no agreement on how close cells need to be in order for them to be considered the same type. Furthermore, even if a distance metric is chosen, for example Euclidean distance in log mRNA fractions, the sparseness and inhomogeneous noise properties of scRNA-seq data make it very challenging to accurately estimate the true distances between cells [12].

In spite of these problems, the current practice in the field is to simply apply *ad hoc* clustering approaches to scRNA-seq data, typically inspired by unsupervised machine learning methods, with the aim of grouping cells of the same ‘type’, e.g. [13, 14, 15]. These clustering approaches generally include several complex layers of data pre-processing, such as normalization and imputation, feature selection, and dimensionality reduction, before the clustering algorithm is applied. These pre-processing steps not only include many fairly arbitrary choices but, as we have recently shown [12], such pre-processing can also severely distort the data by erroneously filtering true biological variability and introducing artefactual correlations. Furthermore, for the clustering itself many different approaches are available, and these typically additionally have many tunable parameters whose values in practice seem to be mostly set by trial-and-error. Given the many layers of *ad hoc* choices involved in these approaches, the resulting clusters lack any biophysical or even methodological interpretation. Instead, the approach taken to confirm the ‘biological validity’ of the clusters, is to show that the cluster exhibit some features that match known biological information, e.g. that certain ‘marker’ genes of a particular cell type are on average higher expressed in a given cluster. However, given that there are combinatorially many different clusterings that exhibit such partial matches with prior biological knowledge, it seems problematic to us to take such partial matches to prior biological knownledge as a validation of the clusters that happened to result from the complex layers of analysis that were applied to the data.

We strongly feel that, instead of applying *ad hoc* clustering methods and attempt to validate these retrospectively by comparison with prior biological knowledge, it is more constructive to first rigorously specify the aims of the analysis, and then *derive* the appropriate algorithm that accomplishes these aims from first principles. This is the approach we take here. We are not going to attempt to solve the general problem of how to define cell types and how to identify them, for reasons laid out above. Instead, our aim is to use clustering so as to maximally reduce the complexity of the dataset without losing *any* of the structure in the data. In particular, we aim to partition the cells of an scRNA-seq dataset into subsets such that the gene expression states of all cells within each subset are statistically indistinguishable. We thus aim to cluster cells at the highest possible level of resolution that is statistically meaningful, i.e. within each cluster all cells are within measurement noise in expression state, and between clusters the expression states are all distinct.

Because the nature of biological and measurement noise in scRNA-seq experiments is known, as characterized in previous studies [16], this task has a uniquely defined solution determined by first principles, as we show below. The resulting method, Cellstates, directly clusters the unnormalized data so that any pre-processing steps are avoided, measurement noise is properly taken into account, and there are no free parameters to tune. For example, the number of clusters is determined by the data, in contrast to most approaches in which the number of clusters is tuned by the user. Moreover, the resulting clusters have a clear and simple interpretation.

Because Cellstates only groups cells whose expression states are statistically indistinguishable, it typically divides the data into many more subsets than other clustering algorithms. To allow comparison with the more coarse clusterings provided by other methods, we additionally provide methods for hierarchically merging Cellstates’s clusters into coarser clusters and to identify marker genes associated with each branching in this hierarchy. As we show below, marker genes of conventionally annotated biological cell types typically correspond to coarser clusters in this hierarchy, allowing us to interpret Cellstates’s clusters as subtypes of conventionally annotated cell types.

## 2 Methods

### 2.1 Multinomial noise in scRNA-seq data implies a parameter-free solution for probabilities of partitions of cells into states

The internal gene expression state (GES) of a cell c, which we will also refer to as a cellstate, is determined by a multitude of biological processes that influence the transcription rates λ_*gc*_(*t*) and degradation rates *μ*_*gc*_(*t*) of mRNAs across genes *g* and time *t* in the history of the cell.

These rates determine the probabilities for the mRNA counts in the cell, which in turn ultimately determine the probabilities of the number of reads captured in a UMI-based scRNA-seq measurement. The probability distribution for the number of mRNAs in a cell *m*_*gc*_ follows a Poisson distribution with mean *a*_*gc*_ given by

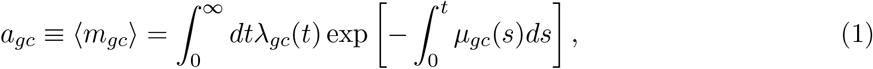

where the time is measured backwards from the present (*t* = 0) to the distant past (*t* = ∞) in the history of the cell [12]. Thus, notably, for each gene *g* in each cell *c*, the entire complex history of transcription rate and mRNA decay rate can be summarized into a single parameter *a*_*gc*_ that fully determines the probability distribution of its current mRNA count for gene g. The scRNA-seq measurement process is noisy and typically only a small fraction (∼ 20% or less) of cellular mRNAs are captured. As this capture rate can vary substantially between cells, information about absolute gene expression levels is lost, at least to some extent. Therefore, more accurate inferences can be made regarding the expected *fractions* of total cellular mRNA that mRNAs of each gene g represent. Following [12], we denote these fractions by transcription quotients α_*gc*_, which we define by

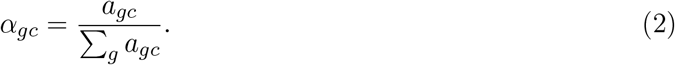

We now define the GES of a cell as the vector 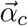 of transcription quotients across all *G* genes. Thus, a GES is a point in the *G*-dimensional simplex α_*gc*_ ≥ 0 ∀g with 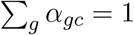.

Given an scRNA-seq dataset with *N* cells, we will assume that the GESs of the cells derive from an unknown set *S* of GESs, where each GES *s* ∈ *S* is characterized by a distinct vector of transcription quotients 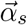. That is, we assume that there are somewhere between 1 (all cells having the same GES) and *N* (all cells having a distinct GES) cellstates represented in the dataset. Our goal is to derive which cells are in the same state and thus separate differences in UMI counts due to biological and measurement noise from differences in the underlying biological state. Thus, the space of hypotheses for this problem is the space of possible partitions of the *N* cells into non-empty non-overlapping subsets. In particular we aim to calculate a likelihood for each possible partition that quantifies how probable the data is under the assumption that all cells in each subset of the partition are in the same GES.

The first step is to derive the relationship between the GES of a cell *c* characterized by 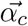 and the vector of its measured UMI counts 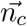, which is summarized in Fig. 1A. Given the transcription activities *a*_*gc*_ of a cell *c*, the mRNA counts are not uniquely determined, but due to inherent biochemical noise in the gene expression process, the mRNA counts *m*_*gc*_ are given by Poisson samples with means *a*_*gc*_. Defining the total transcription activity *A*_*c*_ = Σ_*g*_ *a*_*gc*_, the expected mRNA count for gene *g* can be expressed as the product of *A*_*c*_ and the transcription quotient *α*_*gc*_:

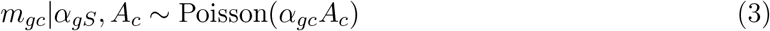

**Figure 1:**
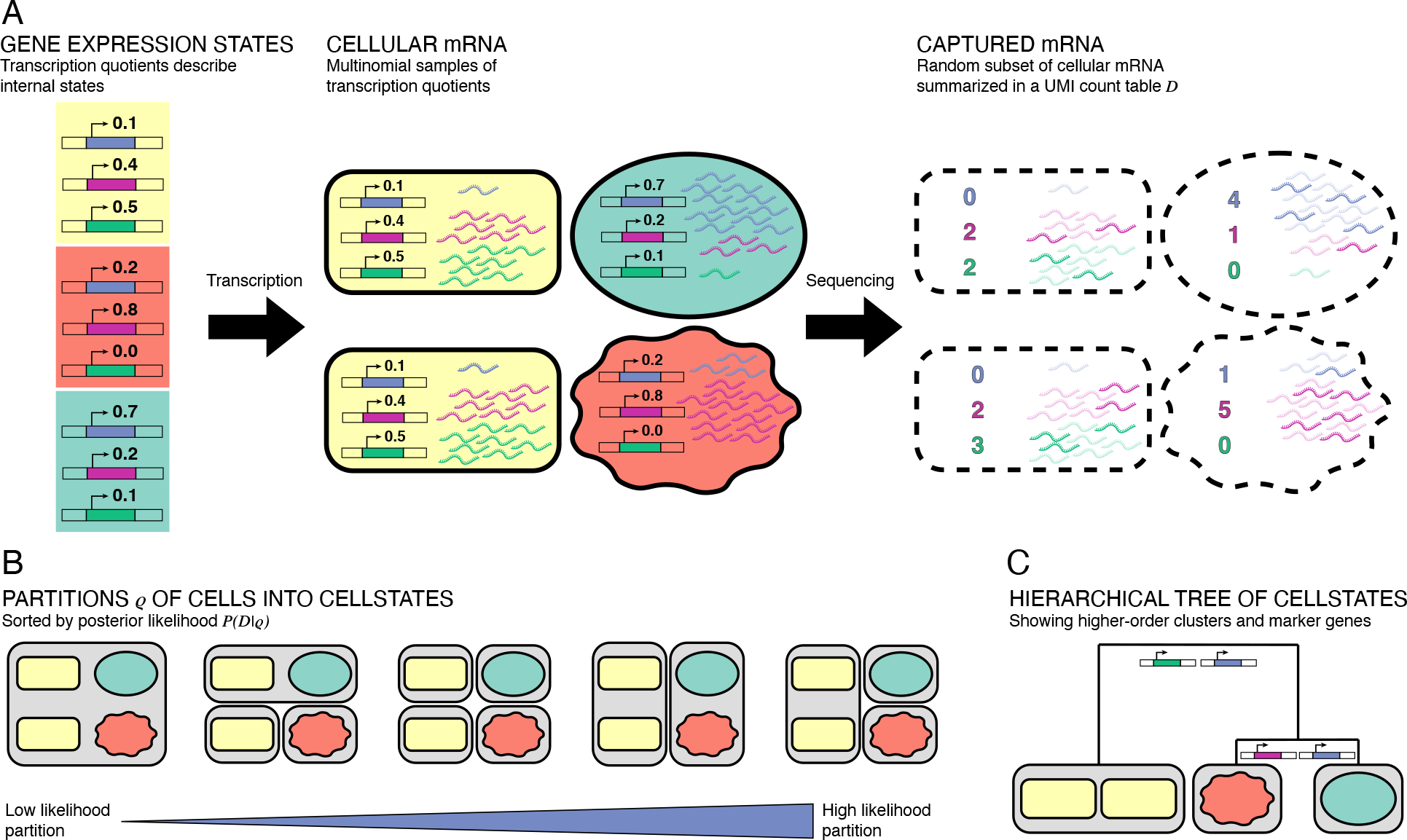
(A) Summary of model assumptions. Each cell is in a gene expression state (indicated by shape and color) characterized by the transcription quotients across genes. The relative numbers of mRNAs in the cell follow a multinomial distribution of these rates. The counts obtained from sequencing reflect a random subset of captured cellular mRNAs and follow the same multinomial. (B) Summary of clustering algorithm. Each partition ρ of cells into clusters gives a likelihood of the data under the noise model. By optimizing the partition, we find groups of cells with shared gene expression states. (C) Cellstates can be hierarchically merged into higher-order cell types. For each merging step, we indicate which genes most contribute to distinguishing the transcription quotients to the left and right below the merger.

Assuming that, for cell *c*, each transcript was captured and sequenced with a probability *p*_*c*_, the distribution of UMI counts *n*_*gc*_ will also be Poisson distributed with mean α_*gc*_*A*_*c*_*p*_*c*_ for each gene *g*. If we marginalize over the unknown capture probability *p*_*c*_ and condition on the total number of mRNAs *N*_*c*_ that were captured for cell *c*, the counts *n*_*gc*_ are simply distributed as a multinomial sample of the transcription quotients 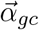 (see the supplementary methods of [12] for a more extensive derivation):

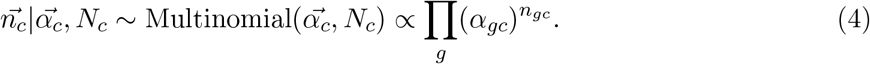

Thus, the probability of the observed mRNA counts 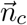 of a cell *c* conditioned on its GES 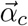 is simply a multinomial sample of size *N*_*c*_ of the expression state 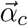. As an aside, we note that the vector of observed UMI counts 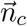 is the unique sufficient statistic for the GES 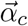 of cell *c*.

Given a partition ρ of the cells into non-overlapping subsets, we now use the above results to calculate a probability *P*(*n* | *ρ*) of the observed UMI counts *n* across all genes and cells, given the assumed partition *ρ*. The derivation of our model follows [17], is explained in detail in the Supplementary Information section A.1.1, and the general approach is illustrated in Fig. 1. Briefly, a partition *ρ* contains subsets of cells s, with one GES 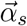 for each subset *s* ∈ *ρ* (i.e. each subset s corresponds to a cluster of cells), and all cells *c* ∈ *s* are assumed to have the same GES 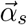 for each subset *s* ∈ *ρ*. The probability for the counts *n*_*gc*_ of all cells in the subset *s* is simply the product over multinomial distributions for each of the cells. Thus, if we define the cluster UMI counts *n*_*gs*_ = Σ_*c*∈*s*_ *n*_*gc*_ and *N*_*s*_ = Σ_*c*∈*s*_ *N*_*c*_, then these cluster counts also simply derive from a multinomial distribution

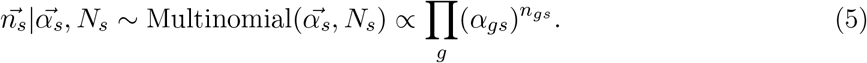

Next, because we do not known the transcription quotients 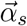 we marginalize over these parameters using a Dirichlet prior. The family of Dirichlet priors is the unique set of priors that is invariant under rescaling of the unknown transcription quotients α_*gc*_ and is parametrized by a vector of concentrations Θ. This marginalization can be done analytically leading to a ratio of products of Gamma functions of the counts *n*_*gs*_ (see Supplementary Information section A.1.1). In this way, a likelihood of the UMI counts 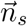 is obtained for each cluster *s* in the partition *ρ*. By taking the product of these likelihoods over all subsets in *ρ*, we arrive at an expression for the likelihood *P*(D|*ρ*, Θ) of the entire dataset 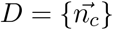 as a function of the partition *ρ* and the parameters of the Dirichlet prior Θ. Taking a uniform prior over both partitions *ρ* and the parameter Θ, the posterior *P*(*ρ*, Θ |D) is simply proportional to the likelihood *P*(D|*ρ*, Θ), which we have obtained in analytical form. The aim of our algorithm is to now find the partition *ρ* and prior parameters Θ that jointly maximize this likelihood.

Importantly, this approach uses only the assumptions that both the inherent biochemical noise in gene expression and the scRNA sequencing introduce Poisson sampling noise, and from first principles derives a parameter-free solution for the most likely partition *ρ*^*\**^ of cells into cellstates that is entirely determined by the raw data *D*. Note that defining cellstates in this way also determines how many distinct cellstates there are, and how many cells there are in each state, directly from the data. The total likelihood of a partition *ρ* simply quantifies how consistent the cells’ measured UMI counts are with the assumption that all cells in each cluster share a common (but unknown) GES 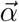. Importantly, over-clustering of the cells into too many cellstates is avoided through the Bayesian framework where increasing the number of GES 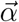 that are marginalized over will lower the likelihood if not supported by the data. To summarize, we partition all cells into subsets such that it is most likely that within each subset the remaining variation between the captured UMI counts is due to random fluctuations.

### 2.2 The likelihood function is optimized using a Markov-Chain Monte-Carlo algorithm

The number of possible partitions of *N* cells grows faster than *e*^*N*^, and we have confirmed that simple greedy searches, such as iteratively fusing clusters of cells to maximally increase the likelihood of the partition, tend to get stuck in local optima of the likelihood function. This makes maximization of the likelihood function challenging. To search for the optimal partition we start from the partition in which each cell forms a cluster by itself and use a stochastic Markov-Chain Monte-Carlo (MCMC) scheme as previously developed in [17]. In each step, a randomly selected cell is proposed to move into a randomly selected different cluster – and accepted if this move increases the likelihood of the partition. If the move decreases the likelihood by a factor *p* < 1, the move is accepted with a probability 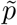 that is adjusted to ensure uniform sampling of partitions (see Supplementary Information section A.2.1). Although theoretically, this MCMC scheme samples partitions in proportion to their likelihood in the long run, we have observed that, in practice, either *p* ≫ 1 or *p* ≪ 1 for most of the proposed moves, most likely due to the fact that the total number of UMI per cell is generally large. Therefore, in practice the optimization essentially performs a random uphill walk to a local optimum rather than sampling the full probability distribution of partitions. We have experimented with a number of different search schemes, including simulated annealing and Gibbs’ sampling schemes, but found that these random uphill walks provide the best balance between total run time and optimality of the final partition. After the MCMC converges, the optimization is followed by some deterministic steps as described in detail in Supplementary Information section A.2.1. Multiple runs of Cellstates on the same data can yield slightly different partitions, and we simply select the best-scoring partition from the partitions obtained in different runs.

### 2.3 Merging cellstates hierarchically into higher-order clusters

As the optimal cellstate partition gives a very fine-grained view of the data, it makes sense to relate the obtained cellstates to each other in a structured manner. To examine the higher-order structure between the cellstates of the optimal partition *ρ*^*\**^, we devised a scheme to hierarchically merge them into higher-order clusters. We define a pairwise cluster similarity as the ratio of the likelihoods of the partition in which the two clusters are merged and the partition in which the two are separated. By construction of the optimal partition *ρ*^*\**^, the similarity will be < 1 for any pair of clusters of this partition and we define a ‘distance’ between two clusters as minus the logarithm of this similarity. The most similar clusters are iteratively merged, resulting in a hierarchical tree of higher-order clusters, see Supplementary Information section A.1.3 and Fig. 1C. As discussed below, we find that these higher-order clusters are often similar to the cell type annotations given in the publications of the datasets on which we ran our algorithm. Additionally, by approximating the multinomial as the product of independent binomials for each gene, we can calculate the contribution of each gene to the cluster similarity score, thus quantifying which genes drive differences in GES between cellstates or higher-order clusters (see Supplementary Information section A.1.4). This allows users of our method to explore the types of cellstates present in the hierarchical tree more easily, i.e. by identifying which genes are associated with particular branchings.

## 3 Results

### 3.1 Cellstates accurately finds optimal partitions in simulated data

As discussed below, we tested Cellstates by running it on a number of published experimental scRNA-seq datasets. However, since there is no ground-truth information available for the GESs of cells in real scRNA-seq datasets, we decided to validate our likelihood maximization algorithm on synthetic datasets that were generated so as to be in agreement with the noise model described above. To get realistic simulated data, we modelled the simulations after results obtained by Cellstates from 18 of the analyzed real experimental datasets as follows. For each of the 18 real datasets we took the optimal partition inferred by Cellstates and then for each of the clusters *s* in this partition, sampled the UMI counts of its cells from a multinomial distribution with mean equal to the inferred GES 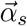 of the cluster. We generated three independent simulated datasets for each of the 18 scRNA-seq datasets, for a total of 54 simulated datasets (See Supplementary Information section A.2.2 for details). We then ran the Cellstates algorithm three times on each simulated dataset. For the large majority of runs Cellstates found the exact same partition as the one that generated the data (Figure S1), which is remarkable since only a tiny fraction of the total space of partitions is sampled during the MCMC likelihood optimization. Moreover, when the partition that Cellstates found differed from the partition that generated the data, this was because Cellstates found a partition with even higher likelihood than the one that generated the data. In fact, for each of the 54 datasets our MCMC likelihood maximization procedure found a partition with likelihood at least as large as the partition that generated the data, and in ≈ 91% of all runs overall (Figure S1).

To compare the similarity of partitions more quantitatively, we will use the two complementary measures of homogeneity and completeness [18] throughout this paper. These measures quantify how much information (as quantified by the Gibbs/Shannon entropy function) a given partition *ρ* contains about a reference partition *ρ*_*f*_ and are both normalized to lie between 0 and 1. If we imagine that we color cells by their cluster in the reference partition *ρ*_*f*_, i.e. so that all cells within each cluster of *ρ*_*f*_ are given the same color, then homogeneity measures how much information the cluster membership in *ρ* provides about the color of the cells (i.e. cluster membership in *ρ*_*f*_). Homogeneity is 1 when, for each cluster of *ρ*, all cells have the same color. Completeness, vice versa, measures how much information the color of a cell provides about its cluster in *ρ*. Completeness is 1 when, for each color from *ρ*_*f*_, all cells of a common color occur in only one cluster of *ρ*. Note that two measures are necessary because if *ρ* is the partition in which each cell is its own cluster, homogeneity is 1 by definition (but completeness is 0). Vice versa, if *ρ* is the partition in which all cells are in one cluster, than completeness is 1 (but homogeneity is 0).

Comparing the partitions inferred by Cellstates on the simulated data to the corresponding reference partitions used to generate the data, we find that they overlap very well, with completeness and homogeneity larger than 0.95 for all runs, and larger than 0.9975 for 118/162 (73%) of the runs, as shown in Figure 2A. As discussed in Supplementary Information section A.2.2, most ‘errors’ occur when the maximum likelihood partition found by Cellstates is higher than that used to generate the data. This happens in particular when the simulated datasets are too noisy to resolve all ground-truth states because the total UMI counts of cells in the ground-truth states are smaller than in the original data.

**Figure 2:**
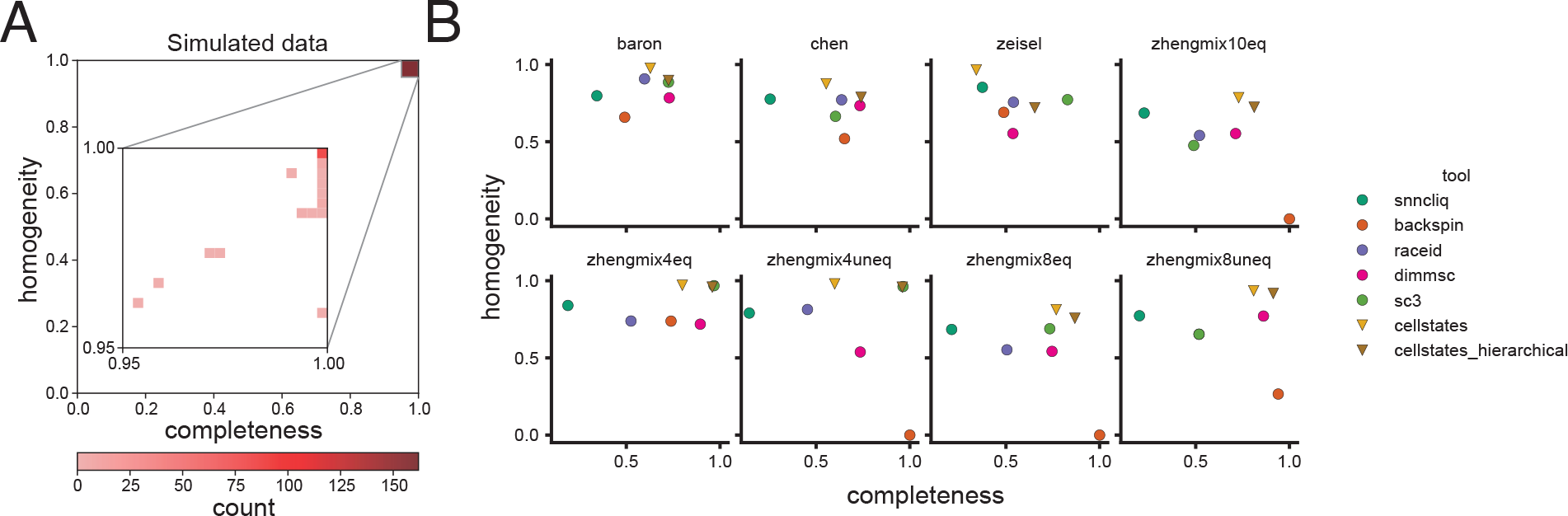
Benchmarking of Cellstates. (A) 2D-histogram of homogeneity and completeness of inferred cellstate memberships in 162 simulated datasets. The inset shows the distributions in the region [0.95, 1], [0.95, 1] where all results fall. (B) Comparison of the performance of Cellstates with those of other clustering tools. For Cellstates, we show the results for the full partition into cellstate-groups (“cellstates”) and merged to the same number of clusters as the annotation (“cellstates_hierarchical”). In each plot, we show homogeneity and completeness of the partitions obtained by the different methods using the published annotation as the reference partitions. Note that low homogeneity and completeness may indicate under-clustering and over-clustering, respectively.

In summary, our tests with simulated datasets show that on datasets that mimic real data, Cellstates performs extremely well on recovering the ground truth used to generate the data, most often recovering the exact partition. And when there is a difference in the partition found, this is most often because Cellstates found an even better partition, which is always very close to the ground-truth partition, as measured by completeness and homogeneity.

### 3.2 Cellstates yields highly reproducible partitions on real datasets

We gathered a total of 29 published datasets from UMI-based scRNA-seq experiments, covering a large range of experimental protocols, tissues and two species (mouse and human), as summarized in Supplementary Table S1. We ran Cellstates on five times on all datasets and compared the best-scoring partition from the five runs with the partitions from the other four runs. We find that the agreement between multiple runs of Cellstates is high, with 88% of the homogeneity and completeness scores larger than 0.9 (Figure S3). These results show that, even though different runs yield different partitions, they do not change substantially between runs.

### 3.3 Cellstates partitions agree better with published annotations than those of other clustering tools

We next compared Cellstates partitions on real datasets with those of a set of previously published methods (BACKSPIN [19], DIMM-SC [20], RaceID [21] and SC3 [22]). A short summary of these methods is provided in Table S2. Assessing the relative performance of different clustering algorithms on real datasets is challenging because in general the ground truth is not known. Here we consider two tests. First, we selected 3 scRNA-seq datasets for which hand-curated annotations of cell types were provided in the publication [23, 24, 19] and compared the partitions obtained by each of the clustering methods with the published annotation. Although there is of course no guarantee that the published annotations are correct, it is reasonable to assume that a better match with these published annotations generally indicates better performance. Second, we also generated a set of five *in silico* mixtures of pure cell populations with known identity, as has been done before for similar benchmarks [25, 16], using data from [26], and checked to what extent each algorithm can correctly recover these known mixtures.

We ran each of the methods on each of these 8 datasets, without tweaking any of the default parameters. Only when a method required the cluster number to be set, we set it to the correct number of annotated cell types. For Cellstates we obtained both the partition given by the method without specifying any parameters, and the partition obtained when hierarchically merging clusters until the number of clusters matches the number of annotated clusters. As for our tests with the simulated data above, we compared the clusters obtained by each of the methods to the annotated clusters through homogeneity and completeness (Figure 2B).

Notably, the partitions found by Cellstates always had the highest homogeneity. This supports that Cellstates indeed only clusters together cells that are in the same underlying gene regulatory state and that this works automatically without the need to correctly set model parameters. However, Cellstates typically partitions the data in more clusters than the annotation, so that the completeness is generally significantly below one. When cellstates are hierarchically merged until the number of clusters matches the annotation (cellstates_hierarchical in Figure 2B), the completeness increases substantially, typically without lowering homogeneity by much.

On the three datasets with published annotations, Cellstates obtained partitions that match the published annotations well, especially in comparison to the partitions produced by the other methods, with only SC3 showing a similar performance on these annotated datasets. Importantly, for the *in silico* cell mixtures – which are the closest we have to a ground-truth annotation – Cellstates clearly outperforms the other methods, and often by a substantial margin. Overall, these results suggest that Cellstates can correctly predict higher-order cell types in scRNA-seq data and that it can do so more accurately than other tools. Moreover, this performance is obtained without any need to pre-process the data or any adjustable model parameters.

### 3.4 Cellstate diversity patterns depend on tissue of origin and not on technical features of the experiment

We next investigated the variation in the number and sizes of the clusters that Cellstates infers on different real datasets. Because measures such as the absolute number of cells per cluster obviously scale with the total number of cells sequenced, we decided to focus on the distribution of cellstate abundances *f*_cellstate_, i.e. the fraction of cells associated with each cellstate. The distribution of *f*_cellstate_ reflects the diversity of different GESs present in a given dataset. As an example, Figure 3A shows the distribution of *f*_cellstate_ for the data from the mouse cortex and hippocampus of [19]. As illustrated by Fig. 3A, we find that *f*_cellstate_ typically varies over multiple orders of magnitude, and that a substantial fraction of the clusters correspond to singlets, i.e. where GESs were only associated with a single cell. That is, the counts in these cells are statistically different from those of all other cells. To obtain a quantitative measure of diversity we looked at various statistics of the distribution of *f*_cellstate_ including the fraction of cells that are singlets, the average cellstate abundance ⟨*f*_cellstate_⟩, its median, and the entropy of the distribution of *f*_cellstate_.

**Figure 3:**
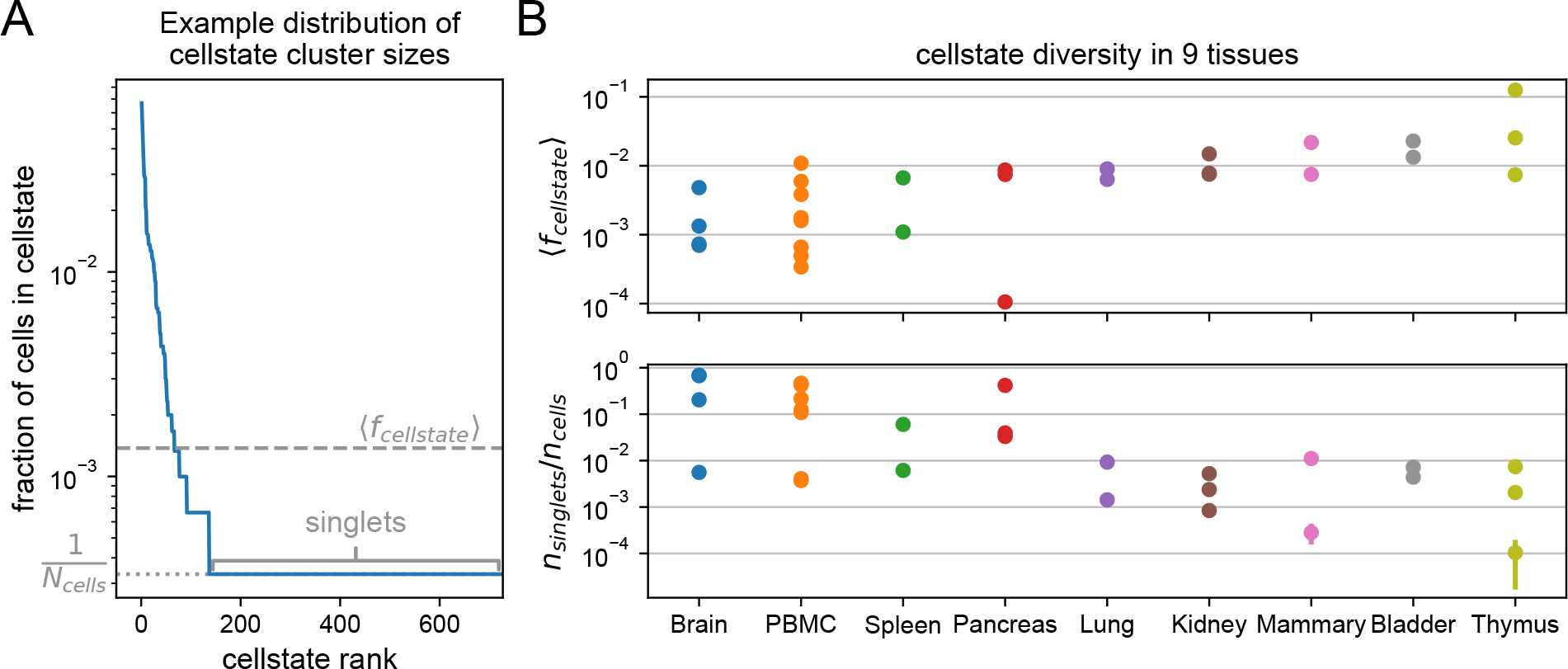
Cellstate diversity reflects the tissue of origin of the data. (A) Example rank-abundance curve for the fraction of cells associated with each cellstate in the dataset from [19]. Such curves describe the diversity of gene expression states in a dataset. The length of the horizontal tail gives the number of singlet cellstates *n*_singlets_ with only one cell; the average cellstate abundance ⟨*f*_cellstates_⟩ is also annotated. (B) For each of the Cellstates results for 29 different datasets from 9 tissues the average abundance ⟨*f*_cellstates_⟩ and the fraction of singlets *n*_singlets_/*n*_cells_ are plotted by the tissue they originate from. These diversity measures show a clear dependence on the tissue, despite the large variation in experimental set-ups used. Error bars show the standard deviations from 5 independent runs of Cellstates and are often so small that they are not visible.

Although ideally these diversity measures would directly reflect the underlying biology of the tissue from which the data derives, we expected that these diversity measures might also strongly depend on technical features of the experiment such as the total number of cells and the typical total number of UMI per cell. For example, the higher the total UMI count per cell, the easier it becomes to distinguish subtly different cellstates, so that one would expect the cellstate diversity to increase with total UMI counts. Similarly, one would expect that the more cells are sequenced, singlet clusters should become less common. To investigate systematically to what extent the distributions of *f*_cellstate_ reflect underlying biology versus technical features, we collected 29 scRNA-seq datasets from 9 different tissues, from different labs and using different sequencing technologies (see Table S1), and investigated how the various diversity measures varied across tissues and with technical features such as cell number and total UMI counts.

Remarkably, we find that all these diversity statistics vary over several orders of magnitude across datasets. For example, the fraction of cells that are singlets varies over three orders of magnitude among the analyzed datasets, from 5 × 10^*−*4^ to nearly 7 × 10^*−*1^ (see Figure 3B), and similarly for the other statistics (Fig. 3B and Fig. S4). Moreover, although there is a lot of variation across datasets, Fig. 3B and Fig. S4 also show that all the diversity measures systematically depend on the tissue of origin of the sample, despite vastly different experimental protocols used to obtain and sequence them. For example, for datasets stemming from biologically diverse cell populations such as brain or peripheral blood mononuclear cells (PBMC), Cellstates correctly and automatically infers few cells per GES, whereas vastly lower diversity is inferred for datasets stemming from Thymus. Moreover, and somewhat to our surprise, the diversity measures show almost no correlation with technical features such as total UMI counts and cell number (Figure S5). The only clear correlation observed is a negative correlation between number of cells and median of the cellstate fractions. This correlation is explained by the fact that the median cellstate fraction often corresponds to singlets, i.e. *f*_median_ = 1/*n*_*cells*_.

In summary, we find that the diversity of cellstates that is found in different datasets reflects the underlying biology of the system, and does not systematically depend on technical features of the experiment. Given this, the observation that in most of the datasets analyzed a large fraction of the clusters are singlets, strongly suggests that the true biological diversity of cellstates is still severely under-sampled in these datasets. That is, many more cellstates exist than are captured at these sampling depths.

### 3.5 Cellstates captures diversity of gene expression states in the mouse brain

Finally, to illustrate how the Cellstates data analysis pipeline can be used for in-depth analysis of a given dataset, we focused on the dataset of [19] consisting of 3005 cells from the somatosensory cortex and from the CA1 region of the mouse hippocampus. Cellstates infers a remarkable diversity in this tissue, with a total of 763 different GESs. Almost a quarter of the cells (727/3005) is in a unique singlet state, but there are also GES with up to 201 cells (7% of all cells), as can be seen in Fig. 3A.

Visualizing how this large number of cellstates relates to another is difficult because the GESs are objects in a very high-dimensional space. The approach that is currently by far most popular in the field is to use stochastic embedding methods that attempt to place cells that are close in gene expression space near each other in a 2-dimensional plane, in particular UMAP [27] and t-SNE [28]. However, it is well appreciated that proper application of these tools is challenging [29], and that beyond approximate conservation of close-neighborhood relationships, the large scale structure in these visualizations is virtually meaningless. In fact, we share the opinion of some in the field that the current usage of t-SNE and UMAP visualizations may be doing more harm than good [30]. Nonetheless, since such visualizations have become the *de facto* standard in the field, we decided to illustrate how cells with different cellstates are placed within a t-SNE visualization of the data (Fig. 4A, left panel).

**Figure 4:**
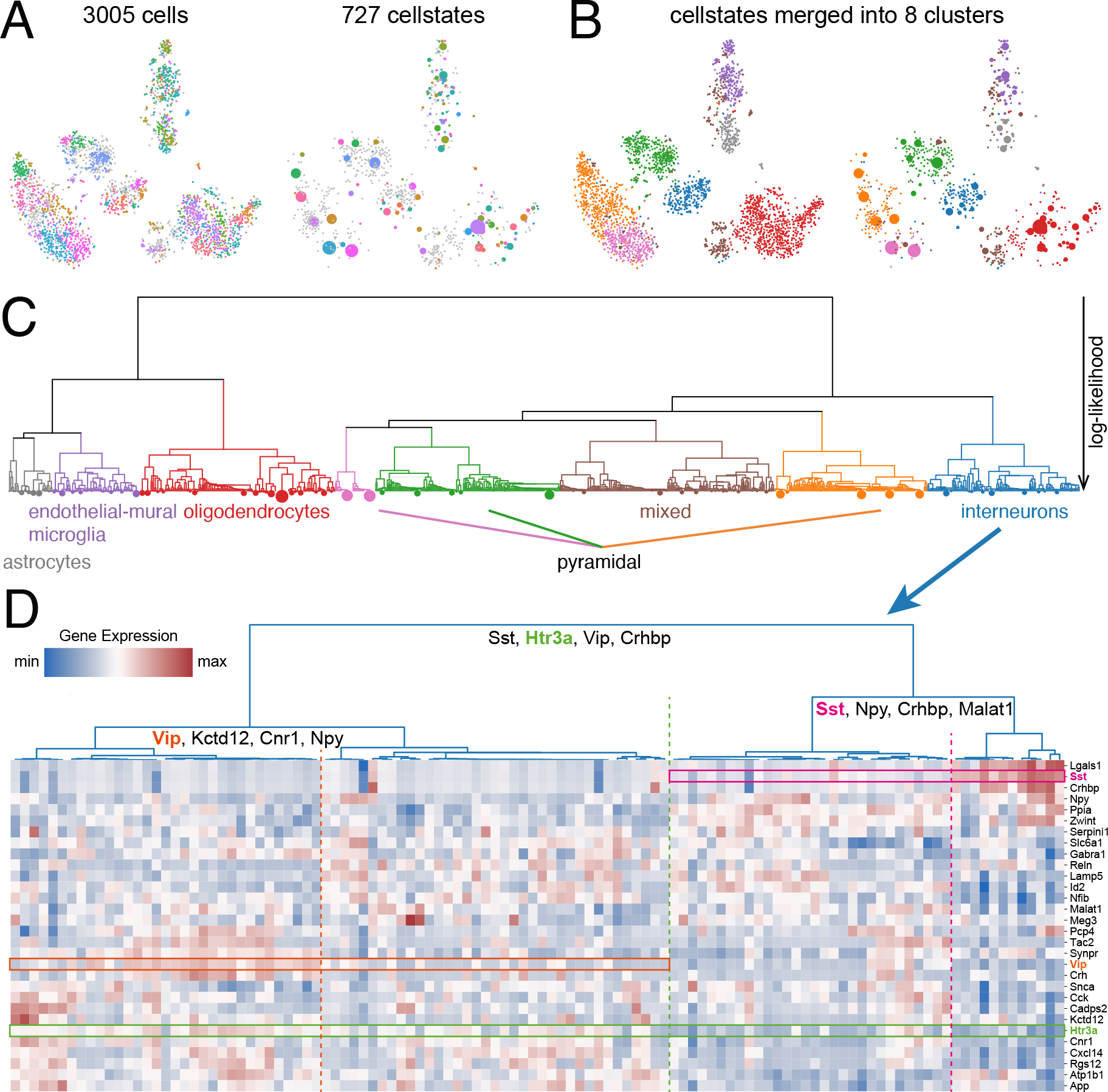
Example analysis of a mouse cortex and hippocampus dataset [19] with Cellstates. (A) Visualization of the data using t-SNE. The colors represent the inferred cellstates, with all singlets shown in gray. One the left, cells are shown individually while on the right cells in the same state were merged into discs. This plot contrasts regions of large gene expression diversity with many small clusters and singlets with low-diversity regions with fewer large clusters. (B) The eight higher-order clusters shown in the same visualization as in (A), with colors representing the eight higher-order clusters defined in (C). (C) Hierarchical higher-order relations between the cellstates. Leaves of the tree correspond to cellstates with their area proportional to the number of cells in them. The vertical height of the branches indicates the negative log-likelihood of the corresponding partition. This tree allows us to split the data into eight higher-order clusters that correspond well to the cell types annotated in [19]. (D) Heat map of gene expression in the interneuron-cluster. Every column corresponds to one GES and shows the corresponding expression pattern. The hierarchical tree shown on the top corresponds to the interneuron sub-tree of (C). Rows correspond to selected genes that are predicted to be differentially expressed between these GESs. In particular, for the first three splits in the tree, top genes contributing to their separation are indicated. Three of these are highlighted in the heat map in green, cyclamen, and orange.

This visualization confirms that cells that are predicted by Cellstates to have the same underlying GES (indicated by the marker color, with singlets in gray) tend to be placed more closely in the t-SNE visualization. Cellstates infers that any variation between cells in the same GES is due to noise, which argues that we can collapse cells in each cellstate and replace them with a single disc at the average of their positions in the t-SNE visualization (right panel in Fig. 4A, with the area of the disc corresponding to the number of cells in the cellstate). This illustrates Cellstates’ ability to reduce the complexity in the data, allowing for a tidier visualization which, for example, highlights that different common cellstates (large colored discs) have different numbers of singlets (gray dots) in their neighborhood.

Next, we hierarchically merged the cellstates to determine higher-order clusters in the dataset, and again visualized the results by coloring either the cells or cellstates in the t-SNE visualization (Fig. 4B). We see that when cellstates are merged into 8 higher-order clusters these clusters largely match the structure of the t-SNE visualization.

However, we feel that a more useful visualization of the relations between the cellstates is obtained by displaying the hierarchical tree resulting from iteratively merging the statistically most similar clusters (Fig. 4C). The tree indicates which cellstates and higher-order clusters are most similar in expression, although it should be remembered that ‘distance’ between clusters is here measured in terms of how statistically significant the differences in the expression patterns are, as opposed to in terms of the magnitude of the changes in gene expression. Notably, at 8 higher-order clusters we find good correspondence with the cell type annotation given in the original publication (Fig. S6), and this is also confirmed by expression of marker genes for these annotated cell types (Fig. S8).

There are however two main differences. Firstly, at this level of resolution in our cluster hierarchy, the annotated clusters of endothelial-mural cells and microglia are merged and they separate only at 15 higher-order clusters (Fig. S7). Secondly, one cluster had a mixed annotation at the chosen resolution. As shown in Fig. S9, this cluster contains cells that express genes which are considered markers for multiple different cell types. This indicates that either the assumption that these genes are markers for specific cell types is incorrect or, alternatively, a technical artefact in the data, e.g. these ‘cells’ might correspond to multiple cells getting the same cell-barcode and having their counts combined.

Finally, we also provide software to extract genes that contribute most to separating the GESs on opposite sides of each branch. To illustrate the use of this tool we focus on the cluster of interneurons, which was particularly diverse with 98 out of 290 cells in singlet states. For each node in the subtree corresponding to the interneurons we identified the genes that contribute most to the separation between the cellstates at opposite branches below it (Supplementary Information section A.1.4), and in Fig. 4D plotted the expression of these genes in a heat map with rows corresponding to genes and columns to individual cellstates. As expected, all columns display unique gene expression patterns, confirming that there are clear differences between the GESs of all cellstates. Furthermore, for three nodes we highlight one example gene whose expression is clearly distinct between the corresponding branches by a rectangle in the heatmap, with the dotted line separating the cellstates on opposite sides of the branch. It should be noted that the most significant genes are those for which the *average* expression on opposite sides of the branch is most significantly different, but the expression of these genes might be quite variable across the cellstates below each branch.

At the highest level the gene Htr3a (green box in Fig. 4D) contributes significantly to separating the expression of interneuron cellstates to the left and right of the branch (dotted green line in Fig. 4D). Similarly, the genes SSt (cyclamen box and dotted line) and Vip (orange box and dotted line) separate cellstates at branches lower in the tree of interneuron cellstate clusters. In general, we find many known markers of interneuron subtypes among the list of differentially expressed genes including *Sst, Npy, Crhbp, Cnr1, Cck* and *Vip* [31, 32, *19, 33]* which supports the biological relevance of the cellstates that we identified.

These results illustrate how Cellstates uncovers substantial sub-structure among a cells of the interneuron type, with the tree structure illustrating the relationships between these subtypes, and the lists of top differentially expressed genes for each branch in the tree providing information regarding the biological differences between these subtypes.

## 4 Discussion

With the popularization of scRNA-seq a vast number of techniques for normalizing and post-processing single-cell gene expression profiles have been developed, including a large number of methods for clustering cells into ‘cell types’, e.g. reviewed in [34, 35, 25, 13]). These methods typically involve several complex layers of analysis steps including the normalization of the raw data, transformation to logarithmic expression (or other ‘variance stabilization’ procedures), selection of ‘features’ to be used in the analysis, mapping of the data to a lower-dimensional representation, often involving abstract latent spaces, selection of a similarity or distance metric, and selection of the final clustering algorithm by which cells are grouped. Moreover, each of these analysis steps typically comes with tunable parameters.

Our impression of the current practice in the field is that the analysis methods are being deployed almost in a trial-and-error manner, i.e. with researchers iteratively trying out different methodologies and tweaking of their tunable parameters, comparing the results with expectations from prior biological knowledge, until results are obtained that look consistent with prior knowledge and, ideally, make some new suggestions that appear biologically plausible.

We believe this kind of approach to analyzing complex data is extremely problematic. The layers of *ad hoc* processing steps and tweaking of parameters make it virtually impossible to give any unambiguous interpretation of the results, to rigorously compare results across different studies, and prohibit direct comparison of the results with those from other experimental approaches. Instead of iterative trial-and-error tweaking of several layers of *ad hoc* methods, we feel that the proper approach to data analysis is to specify the goals of the analysis and the assumptions about the data with enough precision, such that the proper analysis method is unique and transparently follows from these specifications.

This is the approach we have taken in this paper. Instead of attempting to solve the general and very difficult problem of how to determine which cells belong to the same ‘type’, we aimed to solve the simpler problem of maximally reducing the complexity of a given scRNA-seq dataset without *any* loss of structure, by grouping together all cells whose gene expression states are statistically indistinguishable. We have shown that, once a rigorous specification of the measurement noise relating the gene expression states of the cells to the raw scRNA-seq data is given, the appropriate clustering algorithm solving this problem is uniquely determined from first principles. We derived analytical expressions for the posterior probabilities for partitions of the cells into non-overlapping subsets in terms of the raw UMI counts across all genes and cells in the dataset, without any tunable parameters. Moreover, the clusters in the partition with maximal posterior probability have a clear and unambiguous interpretation: they are the optimal way of splitting cells into subsets with transcriptional profiles that are identical up to measurement noise.

A key assumption that our Cellstates algorithm makes is that the cells in a given dataset derive from a finite number of distinct gene expression states, i.e. that in general there will be groups of cells in identical gene expression states, and one might wonder how realistic this assumption is. It is certainly possible that, rather than a discrete set of states, cells could derive from some continuous manifold of gene expression states and it is interesting to ask whether this would manifest itself in the results that we observed with Cellstates. If cells would derive from a continuum, then no two cells would ever be in the exact same state, and one would expect the number of cellstates to grow systematically as the number of cells increases. However, as we have seen in Fig. S5, we find that the observed cellstate diversity depends on the tissue of origin, and does not show systematic correlations with either number of cells or sequencing depth (i.e. total UMI count per cell). Although these are only fragmentary observations at this point and more in-depth study of this question is required, it hints that perhaps discrete cellstates do exist. However, it should also be noted that most datasets analyzed here have a large fraction of singlet cellstates, i.e. clusters with only a single cell per cluster. This suggests that, for these datasets, we are still largely undersampling the true diversity in cellstates that exist in most tissues and it is conceivable that for some tissues we might observe that the number of cellstates does continue to grow as the total number of cells increases, which might then point to the existence of continuous manifolds of cellstates.

One may also ask to what extent Cellstates is vulnerable to batch effects. Our measurement model makes some simplifying assumptions, such as ignoring potential systematic biases that cause transcripts of different genes to be captured with varying efficiency. We note that such gene-dependent capture efficiencies would not effect the distribution over partitions and the optimal partition *ρ*^*\**^, as long as these capture biases are equal in all cells. In fact, as long as systematic biases are the same for the cells within each cellstate, the optimal partition would even remain the same if cells in different cellstates had different capture biases. However, all analysis of gene expression differences between different cellstates of course do rely on the assumption that capture biases are the same across all cells in a given dataset.

One limitation of our approach is that, since the number of partitions increases faster than exponentially with the number of cells and is vast for any realistic scRNA-seq dataset, there is no way to guarantee that our algorithm finds the global optimum even after re-running the program several times. However, our analyses of synthetic datasets with realistic size and structure shows that, in many cases, Cellstates manages to recover the single exact partition that generated the data, and when the generating partition was not recovered exactly, most often this was because a slightly different partition with even higher likelihood was found (Fig. S1). On real data we also found that the partitions obtained in different runs of cellstates are generally very similar (Fig. S3). These results suggest that the vast space of partitions can be effectively searched by Cellstates’s Monte Carlo Markov Chain procedure.

However, it should be noted that, especially compared to most methods currently used in the field, on larger datasets Cellstates can have long run-times and requires significant computational resources. In the future we intend to improve the speed of the method by using computationally less expensive methods to either first subdivide larger datasets into coarse subsets before running Cellstates or to pre-select neighborhood relationships, i.e. which pairs of cells are candidates for mergers with each other. Nonetheless, we note that although it may take quite some time to run Cellstates on a dataset, it generally is still considerably less than the time required to perform the experiment. Moreover, since Cellstates has no tunable parameters, the method has to only be applied once. In fact, we believe that the strong importance that is currently assigned in the field to having fast analysis methods derives largely from the fact that most researchers apply these methods in a trial-and-error manner, running many times with different parameters settings and filters until results are obtained that ‘look best’ by some preconceived notions of what the data should show. As we already discussed above, we think this ‘fast analysis’ methodology is scientifically unhealthy, and like the movement advancing ‘slow food’ over ‘fast food’, we propose that analysis of complex large-scale datasets in biology would strongly benefit from a ‘slow analysis’ movement that favors slow but rigorously motivated methods over iterative tuning of fast *ad hoc* methods.

Finally, we would like to comment on the way we imagine Cellstates can be applied in practice. The most obvious application of Cellstates, and the one we highlighted here, is to identify subtle substructure among known cell types, the relationships between these subtypes, and the genes that most distinguish these subtypes. However, we feel that an arguable even more important use of cellstates is as a way to significantly reduce the complexity of a dataset without losing *any* structure in the data. That is, after cells have been clustered into cellstates, one can decide to simply treat these clusters as if they were ‘super cells’ and perform further analysis and processing such as trajectory reconstruction, pseudo-time analysis, visualizations or differential gene expression inference treating these clusters as if they were single cells. A recent study has shown that such an approach can indeed lead to improvements in downstream analyses [37], even in the absence of a rigorous methodology for clustering. Cellstates provides precisely the rigorous methodology for reducing the complexity of the dataset and removing some of the inherent noise in scRNA-seq data, while leaving all underlying biological variation completely intact. We propose that this application of Cellstates is an ideal first step in any scRNA-seq data analysis pipeline.

### 4.1 Software availability

The Cellstates python package is available online on GitHub (https://github.com/nimwegenLab/cellstates). It can be run through the command-line on files containing the table of unnormalized expression data (UMI counts) in one of several formats including compressed and uncompressed tab-separated values, Matrix Market, and NumPy binary. The outputs of this command-line tool include the optimized partition of cells into cellstates, the optimized prior parameter Θ, the hierarchical tree of cellstates that can be used to find higher-order clusters, and a table of differential expression scores for each gene and node in the tree. Furthermore, we provide python functions for analyzing these Cellstates outputs, including finding the mean and modal vector of transcription quotients of each cellstate, and visualizations of the hierarchical trees as shown in this paper. We also provide several notebooks with example analyses.

## 5 Acknowledgments

We thank Thomas Sakoparnig for helpful discussions, testing of the code, and for contributing example notebooks with cellstates analysis. We thank van Nimwegen lab members for comments on the manuscript and Daan de Groot for identifying bugs in the code.

## 6 Tables

- S1: Table of datasets.
- S2: Table of clustering tools.

## A Supplementary Information

### A.1 Detailed Derivations

#### A.1.1 Likelihood of partitions

We consider an scRNA-seq dataset D characterized by a matrix of UMI counts *n*_*gc*_ corresponding to the number of UMIs for gene *g* in cell *c*. We denote by *ρ* partitions of the cells into non-overlapping subsets and want to determine a likelihood function *P*(D |*ρ*) under the assumption that, for each subset *s* ∈ *ρ* of the partition, all cells *c* ∈ *s* have the same gene expression state. To explain how this likelihood function is calculated, we first discuss how the mRNA counts *m*_*g*_ of a cell, and observed UMI counts *n*_*g*_ in an scRNA-seq experiment depend on the gene expression processes in the recent history of the cell, as previously introduced in [12].

Given the inherent thermodynamic fluctuations affecting the molecules inside the cell, and the Brownian motion that they are subject to, even a comprehensive description of the current ‘state’ of the cell in terms of the number of molecules of each type in each cellular compartment only determines the *rates* with which different molecular reactions occur. For the mRNA levels of a given gene *g*, the relevant rates are the transcription rate λ_*g*_ and the mRNA decay-rate *μ*_*g*_. It is well established that for a gene g with constant transcription rate λ_*g*_ and constant mRNA decay-rate *μ*_*g*_, the number of mRNA molecules in the cell *m*_*g*_ follows a Poisson distribution with mean ⟨*m*_*g*_⟩ = *a*_*g*_ = λ_*g*_/*μ*_*g*_, e.g. [38]. More generally, when *μ*_*g*_ and λ_*g*_ are arbitrary time-dependent functions *μ*_*g*_(*t*), λ_*g*_(*t*), with λ_*g*_(*t*) denoting the transcription rate a time *t* in the past of the cell, and *μ*_*g*_(*t*) the decay rate of mRNAs for gene *g* a time *t* in the past, the probability distribution for the current number of mRNAs *m*_*g*_ in the cell is still a Poisson distribution with mean [12]:

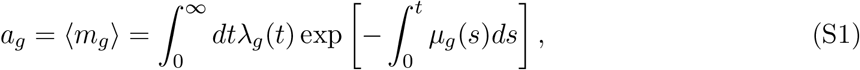

which we call the ‘transcription activity’ of gene g. Note that time is measured backwards from the present (*t* = 0) to the distant past (*t* = ∞) in the history of the cell. Thus, a single parameter *a*_*g*_ for each gene *g* is sufficient to fully characterize the distribution of mRNA numbers in a cell at any given time point. The remaining uncertainty about the actual numbers is due to random thermodynamic fluctuations in events such as RNA polymerase binding or mRNA degradation. To conclude, given the expression state of the cell as defined by the vector of transcription activities 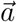, the probability of a count vector of cellular mRNAs across all genes 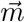 is therefore a product of Poisson distributions:

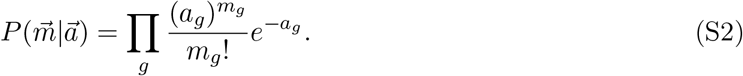

We will assume that the *measured* UMI counts *n*_*g*_ correspond to a random sample of the cell’s total mRNA pool *m*_*g*_ with some unknown capture rate *p* per mRNA. As will be discussed below, our model remains valid if the capture rate *p* varies between cells or has gene-dependent biases. Given these assumptions, the likelihood for the observed UMI count vector 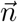 is still a Poisson distribution, albeit with a different mean:

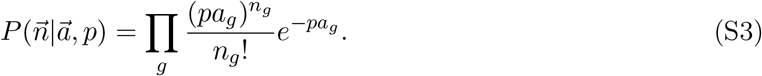

Following [12], we now define the transcription quotients α_*g*_ = *a*_*g*_/*A*, with *A* = Σ_*g*_ *a*_*g*_ the total transcription activity of the cell. Note that α_*g*_ corresponds to the expected fraction of transcripts from gene *g* among all transcripts in the cell. For a cell with a total UMI count *N* = Σ_*g*_ *n*_*g*_, we have

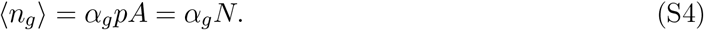

Conditioned on the total count *N*, the distribution of all measured counts 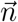 is a multinomial in the transcription quotients:

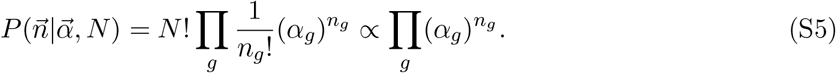

This is the form of the likelihood of the UMI counts of a single cell as a function of the transcription quotient vector 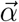.

In our model, we use the transcription quotient vector 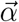 to represent the ‘expression state’ of a cell and the key ingredient of our model is that, given a partition *ρ*, all cells within each subset *s* of the partition have the same transcription quotient vector 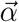. The likelihood for the counts *D*_*s*_ of a subset of cells s that have equal transcription quotients 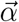, is given by

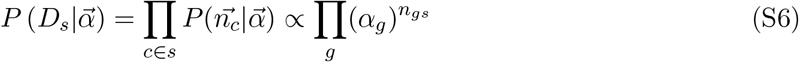

where 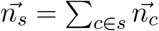 is the vector of total UMI counts among all cells in the subset *s*.

To calculate the likelihood *P*(*D*|ρ) of a partition, we need to marginalize over the unknown transcription quotient vector 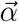 for each of the subsets *s* in *ρ* and to do this we have to define a *prior* distribution over possible transcription quotient vectors 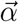. We will use a Dirichlet prior, which corresponds to a maximal ignorance prior in the sense that it is the unique prior that is invariant under arbitrary rescaling of the transcription quotients α_*g*_ → λ_*g*_α_*g*_. Moreover, it is the conjugate prior to the multinomial distribution, allowing us to analytically marginalize over the 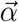. In particular, for each subset s we characterize our prior information regarding its transcription quotient vector 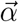 by the *same* Dirichlet prior:

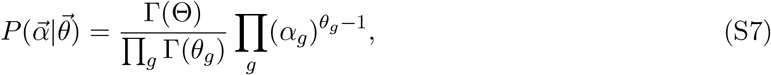

where the θ_*g*_ are the parameters of the Dirichlet prior and Θ = Σ_*g*_ θ_*g*_. We can now marginalize over 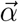 and obtain

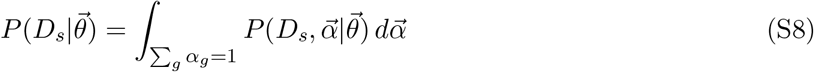

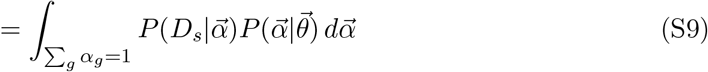

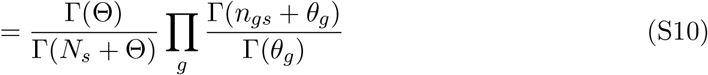

with *N*_*s*_ = Σ_*g*_ *n*_*gs*_ is the total number of UMI summed over all cells in subset *s*. The likelihood of a partition *ρ* is now obtained by simple taking the product of this expression over all subsets *s* ∈ *ρ*:

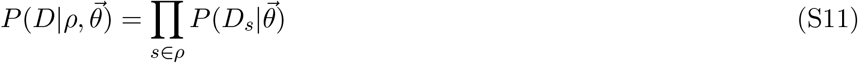

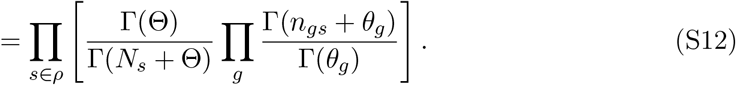

Note that this expression is very similar to likelihood functions derived previously for clustering DNA sequences [17]. Essentially the only change is that the 4-letter DNA alphabet is here replaced by the ‘alphabet’ of *G* genes.

Ideally, we would search for the combination 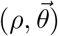 that jointly maximizes the likelihood, i.e. optimizing both the partition *ρ* and the parameters θ_*g*_ for each gene individually, but this is computationally intractable. Without loss of generality, we can rewrite the parameters of the prior θ_*g*_ as the product of an overall scale vector Θ and a normalized vector 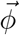 with Σ_*g*_ *ϕ*_*g*_ = 1. Second, we note that for the trivial partition in which all cells are put into a single cluster, the optimal *ϕ*_*g*_ are given by

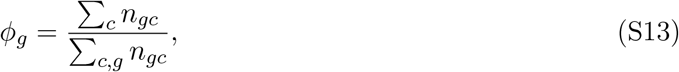

i.e. the prior parameter ϕ_*g*_ simply equals the fraction of UMIs for gene g in the entire dataset. We will simplify the optimization of the prior’s parameters by fixing 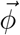 to this vector, setting 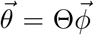, and only optimize the scale factor Θ ∈ ℝ^+^, while leaving the ϕ_*g*_ fixed for a given dataset. Setting the prior in this way ensures that, for each subset s, the expected direction of the transcription quotient vector 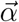 matches the overall UMI counts in the entire dataset, while optimizing Θ allows to tuning of the expected amount of variability around this ‘average’ vector of transcription quotients.

With this chosen form of the prior, we finally get an expression for the likelihood of the whole dataset *D* that only depends on the scale factor Θ and the partition of cells into subsets *ρ*:

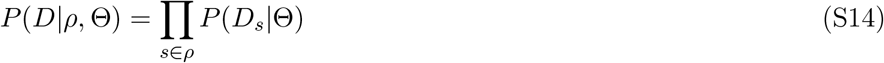

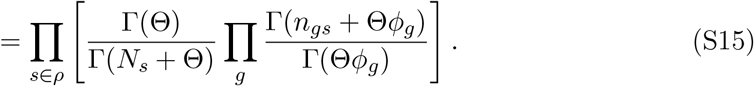

Finally, we return to discuss our simplifying assumption that the mRNA capture rate *p* is constant across genes and cells and show that this assumption can be significantly relaxed without affecting the results. In particular,we can assume that the probability *p*_*gc*_ of capturing (and successfully amplifying and sequencing) an mRNA for gene *g* in cell *c* can be written as

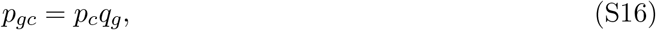

where *p*_*c*_ is a cell-specific overall capture rate and *q*_*g*_ describes gene-dependent biases that may be specific to the particular experiment, but are assumed constant across the cells in the experiment. With this capture efficiency, the expected UMI count for gene *g* in cell *c* is ⟨*n*_*gc*_⟩ = *p*_*c*_*q*_*g*_*a*_*g*_. Thus, the expected fraction of counts from gene *g* becomes

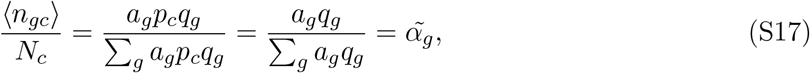

where the last equality defines 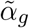. From here, we can proceed the derivation exactly as before with 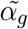 replacing α_*g*_. As we marginalize out this variable in Equation S10, the final result is invariant. Note that, given that we separately marginalize over the 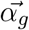 of each subset, the result is even invariant when different subsets *s* have different gene-bias vectors 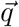, as long as all cells within a subset have the same bias. This suggests that our likelihood over partitions is not only insensitive to fluctuations in overall capture efficiency across cells, but will also be quite robust to fluctuations in gene-dependent capture efficiency as long as cells with equal expression states have equal biases.

#### A.1.2 Posterior of transcription quotients

Although for calculating likelihoods over partitions *ρ*, we marginalize over the GES 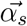 for each subset *s*, we can of course also obtain posterior distributions over these GES for each cluster. Given a subset of cells *s*, the posterior distribution for its GES 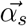 can be obtained from equations S6, S7 and S10 to find:

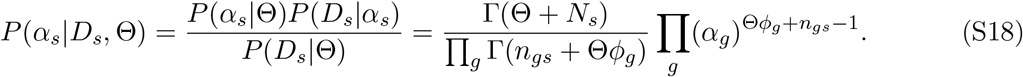

From this expression, we can derive expressions for mode, mean and variance of 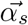:

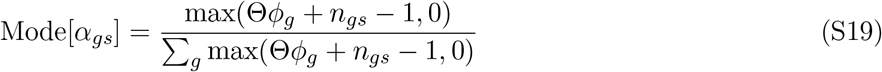

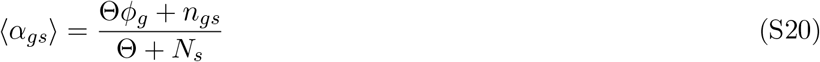

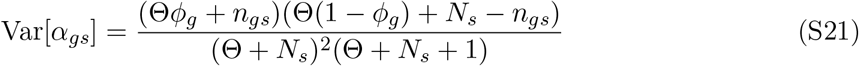

Often we are interested in the log-transcription quotients δ_*gs*_ = log(α_*gs*_) rather than the transcription quotients themselves. For these we find:

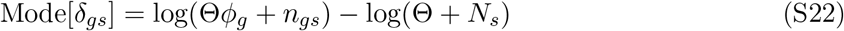

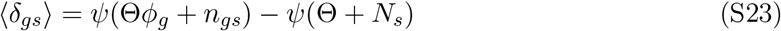

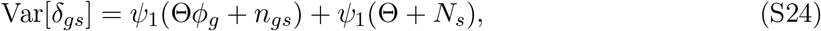

where *ψ* is the digamma function (the derivative of the logarithm of the Gamma function) and *ψ*_1_ is the first derivative of the digamma function.

#### A.1.3 Cellstate similarities and hierarchical clustering

We define the similarity between two cellstates *S*_*a*_ and *S*_*b*_ as the ratio of the likelihoods of the partitions with both subsets merged and each subset separate:

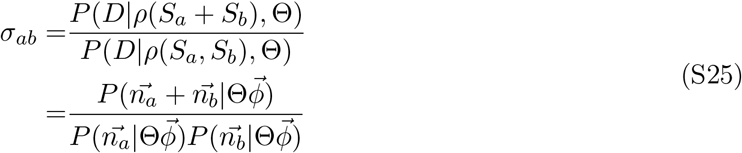

Note that this similarity metric does not behave like usual distance metrics in that similar clusters will have a large *σ* and dissimilar clusters a small *σ* close to 0. To build the hierarchical tree of the cellstates, we start by setting the leaf clusters of the tree to the cellstates of the optimal partition and then calculate all pairwise similarities *σ*_*ab*_ between all pairs of leaf clusters. We then iteratively merge the pair of clusters with the highest similarity, and recalculate the pairwise similarities with the newly formed cluster until all clusters have merged into a single cluster. For plotting, we save the resulting tree in the Newick format with distances set to positive log-similarities d_*ab*_ = max[− log(*σ*_*ab*_), 0].

#### A.1.4 Differentially expressed genes between pairs of clusters

To describe the differences between the cellstates of different clusters, and to help give biological interpretation, it is useful to quantify which genes are most differentially expressed between the clusters. In our framework, we can quite naturally define the extent of differential expression of genes by decomposing Equation S25 into contributions of individual genes, i.e. *σ*_*ab*_ = ∏_*g*_ *σ*_*ab,g*_. A low value of *σ*_*ab,g*_ for a gene g indicates that the differences in UMI counts are much more different between the two clusters a and b than would be expected from noise. A high value *σ*_*ab,g*_, in contrast, indicates that counts are within the expected noise levels.

To obtain such a decomposition, we start by decomposing the cluster likelihood of Equation S10 into contributions from individual genes 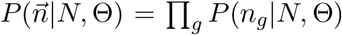. In the multinomial model, the likelihood is conditioned on the total number of captured mRNA in the cell *N*, so that, formally, the counts *n*_*gc*_ are correlated for all pairs of genes. However, since this correlation is generally quite weak, we can make the assumption that the expression noise is independent between genes. Thus,

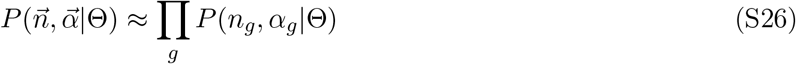

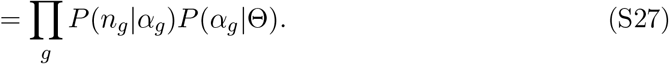

We can find 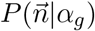 by marginalizing over the other variables:

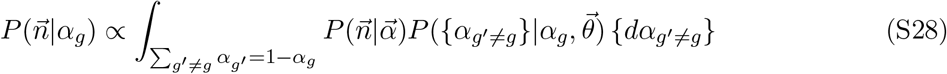

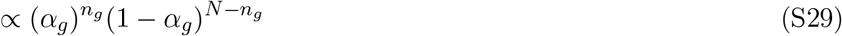

where *N* = Σ_*g*_*n*_*g*_. We can further marginalize over {n_*g′*≠*g*_} requiring that *N* is constant, and normalize to obtain the binomial distribution:

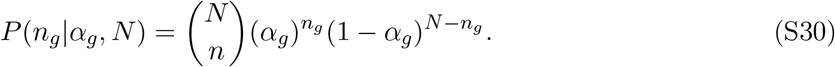

Next, we find P(α_*g*_|Θ):

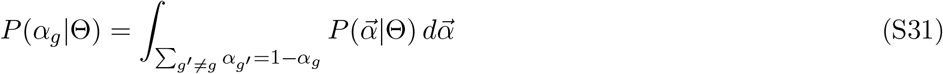

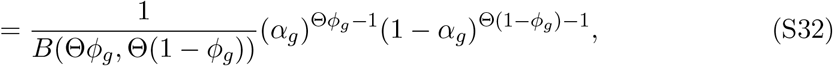

where B(*α, β*) = Γ(*α*)Γ(*β*)/Γ(*αβ*).

Finally, we have all ingredients of Equation S27 above. Unlike in Equation S10 where we perform the integral subject to the constraint Σ_*g*_ *α*_*g*_ = 1, we integrate over all *α*_*g*_ separately. With the high dimensionality of 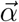 and assuming the likelihood function has a sharp peak, the error will be small.

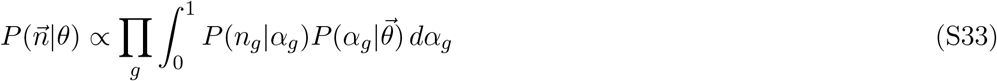

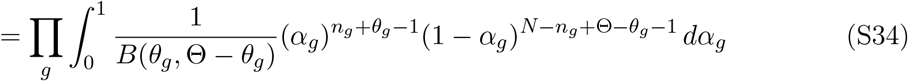

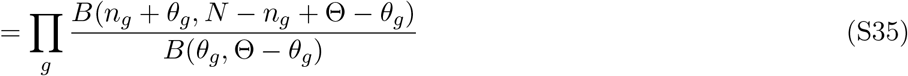

This likelihood function clearly has a separate contribution *P*(*n*_*g*_|*N*, Θ) for each gene:

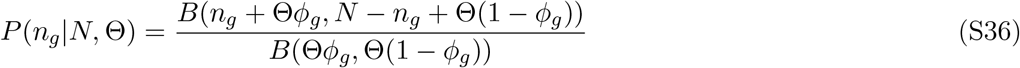

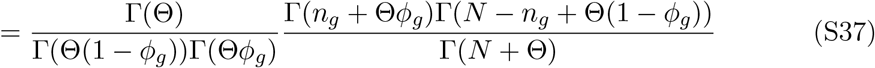

Comparing to Equation S10, we see that this is equivalent to taking the ratio of the cluster likelihood with gene *g* and without *g*.

Finally, we can use this expression for *P*(*n*_*g*_ | *N*, Θ) in Equation S25 and define a gene-specific score for differential expression

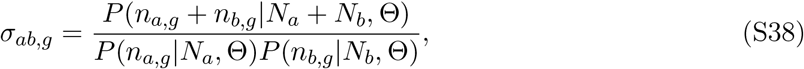

where the subscripts *a, b* refer to two subsets of cells. Genes with *σ*_*ab,g*_ < 1 have counts that poorly fit a model with a single transcription quotient for both clusters, compared to a model where the two clusters have distinct transcription quotients. In contrast, genes with a score *σ*_*ab,g*_ > 1 favour a model with a single transcription quotient and are not differentially expressed between the clusters.

### A.2 Computational Methods

#### A.2.1 MCMC Algorithm for maximizing the likelihood

Our aim is to identify the combination of a scale factor Θ and partition *ρ* that jointly maximize the likelihood *P*(*D* | *ρ*, Θ) given in Equation S14. To do this, we start from an initial guess for Θ and then iteratively

1. Search for the partition *ρ*^*\**^ that maximizes the likelihood with the current value of Θ,
2. Given *ρ*^*\**^, find the value of Θ that maximizes the likelihood *P*(*D*|*ρ*^*\**^, Θ),

until convergence. In order to limit the number of time-costly optimizations of the partition, we only consider values of Θ = 2^*q*^, *q* ∈ ℕ. The initial guess is taken with *q* = ⌊log_2_(⟨*N*_*UMI*_⟩) + 0.5 ⌋ where ⟨*N*_*UMI*_ ⟩ is the average number of total UMI counts per cell. That is, our initial guess for Θ corresponds to the average total UMI count per cell. This means that strength of the influence of the prior is about equal to the influence of the data from a single-cell.

To optimize the partition *ρ* at a given Θ, we start with the partition in which each cell forms its own cluster. To explain how the space of partitions is searched, we conceptualize partitions as putting cells into ‘boxes’, i.e. such that all cells in the same box form a cluster and different boxes correspond to different clusters. Initially we assign each of the *C* cells into one of *C* boxes and we will search the space of partitions by moving cells between these *C* boxes. Specifically, we use a Markov chain Monte Carlo (MCMC) algorithm, which iterates the following steps:

1. Given the current partition *ρ*, a cell is chosen uniformly at random and taken out of its current box.
2. One of the *C* −1 other boxes is chosen uniformly at random and we consider the partition *ρ*^*′*^that is created by moving the cell into this box.
3. We calculate the likelihood ratio of the new to old partitions: *P*_move_ = *P*(*D*|*ρ*^*′*^, Θ)/*P* (*D*|*ρ*, Θ).
4. If the new partition *ρ*^*′*^ has *N*_clus_ clusters and *ρ* had *N*_clus_ +1 clusters, we set *P*_bias_ = (*C* − *N*_clus_), otherwise *P*_bias_ = 1.
5. If *P*_accept_ = *P*_move_ \**P*_bias_ > 1, the move is accepted, otherwise it is accepted with probability *P*_accept_.

Note that, as explained in the supplementary material of [17], the correction factor *P*_bias_ ensures detailed balance, i.e. that in the absence of differences in the likelihoods of the paritions *P*(*D* | *ρ*, Θ), all partitions would be sampled uniformly.

These steps are iterated until the likelihood has stopped increasing for a sufficient number of steps. In the current implementation of the algorithm, the stopping criterion is controlled by two parameters: the number of steps *S* and the number of tries per step *T*. Each round a total number of *S* × *T* moves are attempted. This is repeated if at least *S* of the trials led to an accepted move, i.e. the partition was changed at least *S* times. If less than *S* moves were made in the *S* × *T* trials, the value of *S* is reduced by 10 and the value of *T* is multiplied by 10, and new rounds of trials are started. This is continued until *S* falls below 10, after which the MCMC moves are stopped. Note that the algorithm keeps track of the partition with the highest likelihood it has seen, and set the partition to this highest likelihood partition at the end of the rounds of MCMC moves. By default we set *S* = *N*, i.e. equal to the number of cells, and *T* = 1000, but these values can be changed by the user.

After these MCMC moves, a final uphill walk is performed as follows. For each pair of clusters existing in the partition, we calculate the likelihood change that would occur if the clusters were merged into one. We then iteratively merge clusters until no more mergers are left that would increase the likelihood. Finally, for each cell we calculate the likelihood change that would occur if the cell were moved into any of the currently existing clusters, and move the cell to its optimal cluster.

#### A.2.2 Simulated Datasets

Simulated datasets were created based on the inferred cellstates of real datasets. Of the 36 datasets which were analyzed for this paper, we selected those 18 that have less than 6000 cells, more than 3 cellstates with more than 10 cells in them, and a median number of UMIs per cell greater than 1000. Our aim was to simulate datasets based on the set of transcription quotients inferred to be present in the real datasets. Additionally, we wanted to make sure that the clusters can only be identified by differences in transcription quotients (i.e. relative gene expression levels) and not by differences in total UMI counts. The total number of UMI for each cell *c, N*_*c*_, was therefore drawn independently from a log-normal distribution that was fitted to the experimental distribution of *N*_*c*_ in the corresponding dataset.

For each cellstate *s*, we determined the mean expression transcription quotient vector 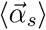 and then sampled the UMI count vectors 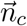 of each cell *c* in the cellstate s from a multinomial distribution with mean 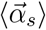. However, we found that the maximal likelihood partition of the simulated dataset often differed from the partition generating the dataset, especially for very small and singlet clusters. In particular, when cells that were in a singlet state were given a lower total UMI count in the simulation, these cells were often no longer statistically significantly different from other states, and this to a lesser extent also affected small clusters. To mitigate this problem, we retained only cellstates with more than 10 cells. The number of cells in each cellstate was kept the same as in the experimental data. With this simulation procedure, we made sure that most simulated cellstates would be statistically distinct, even when total UMI counts were lower than in the original dataset.

In this way, three separate simulated datasets were generated for each experimental dataset. Lastly, an additional three “down-sampled” simulations were carried out with ⟨log_10_(*N*_*c*_)⟩ = 3 and var(log_10_(*N*_*c*_)) = 0.1 fixed for all datasets.

#### A.2.3 Further Discussion of the results on simulated data

In Figure S1 we show, for each of the three simulations (marked by their color) generated per experimental dataset (different columns), the detailed outcomes of three independent Cellstates runs per simulation. The top panel shows the difference in log-likelihood Δ*L* between the inferred partition and the partition used to generate the simulated data. Negative scores mean that the inferred partition had a lower likelihood than the simulated one and can be attributed to a failure of the algorithm to find the optimal partition. As can be seen, such errors are rare and there is always at least one out of the three runs that has Δ*L* ≥ 0, i.e. a partition at least as good as the one used to generate the dataset was always found. Positive scores indicate that a partition was found with a higher likelihood than the one used to generate the dataset. This can happen for example if there are not enough counts in the simulated cells to statistically distinguish cellstates. To test this hypothesis, we looked at the “down-sampled” simulations with fewer UMI per cell which would make cells with similar, but distinct transcription quotients indistinguishable. Indeed, the results shown in Figure S2 confirm this hypothesis: For most down-sampled simulations, the best-scoring partition is different from the ground-truth. Also, they tend to score low on homogeneity but high on completeness - which means that inferred clusters are unions of clusters used to generate the simulated data.

### A.3 Supplementary Figures

**Figure S1:**
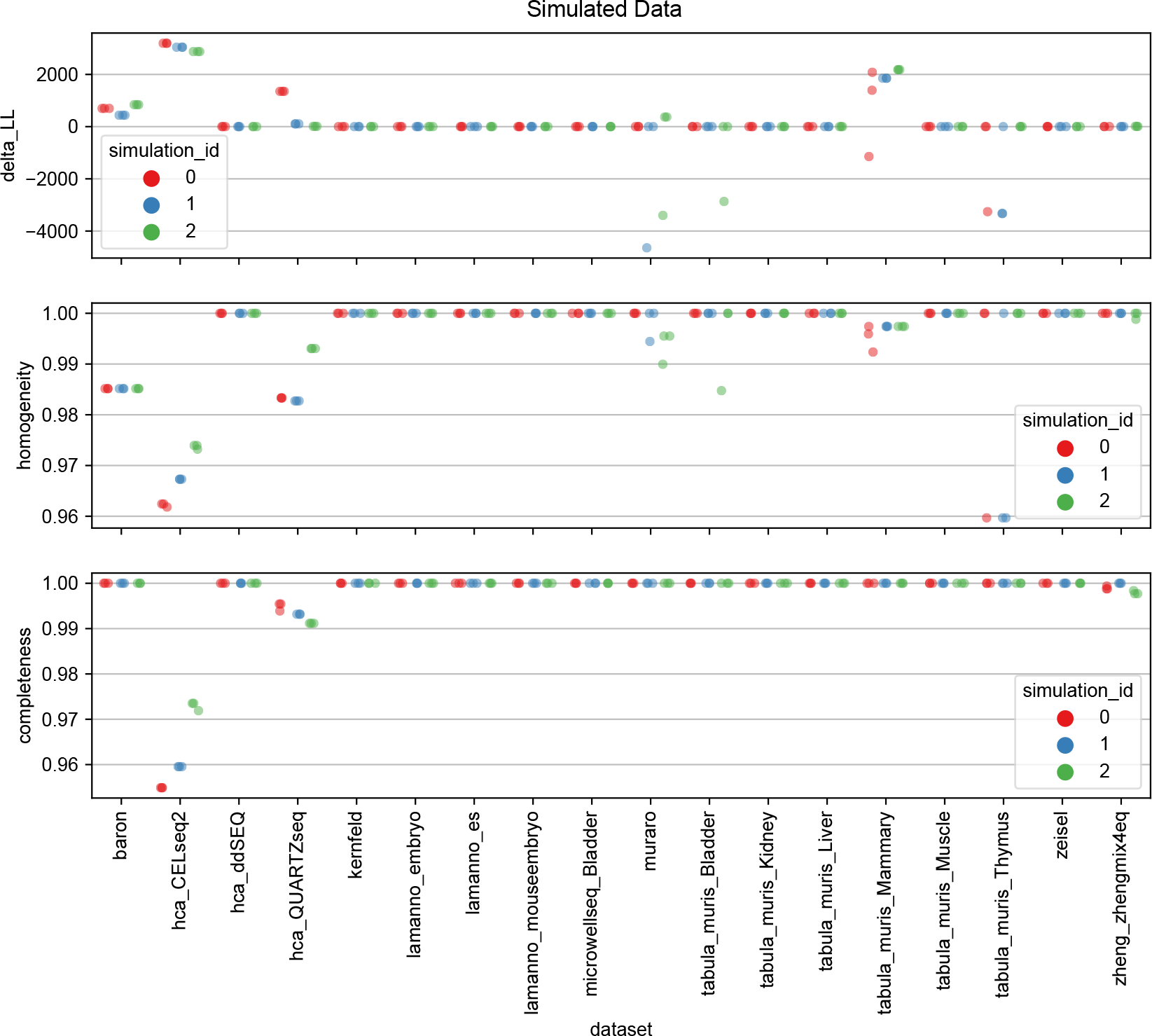
Detailed results from Cellstates runs on simulated data based on the set of GESs from various indicated datasets. For each of the 18 real datasets, three simulations were generated (red, blue, and green) and Cellstates was run three times on each simulated dataset. In the top panel, the difference in log-likelihood delta_LL between the inferred and simulated partitions is shown (with a positive difference meaning that a partition was found with higher log-likelihood than the one used to simulate the data). The corresponding homogeneity and completeness of the inferred compared to the simulated partitions are given in the two lower panels.

**Figure S2:**
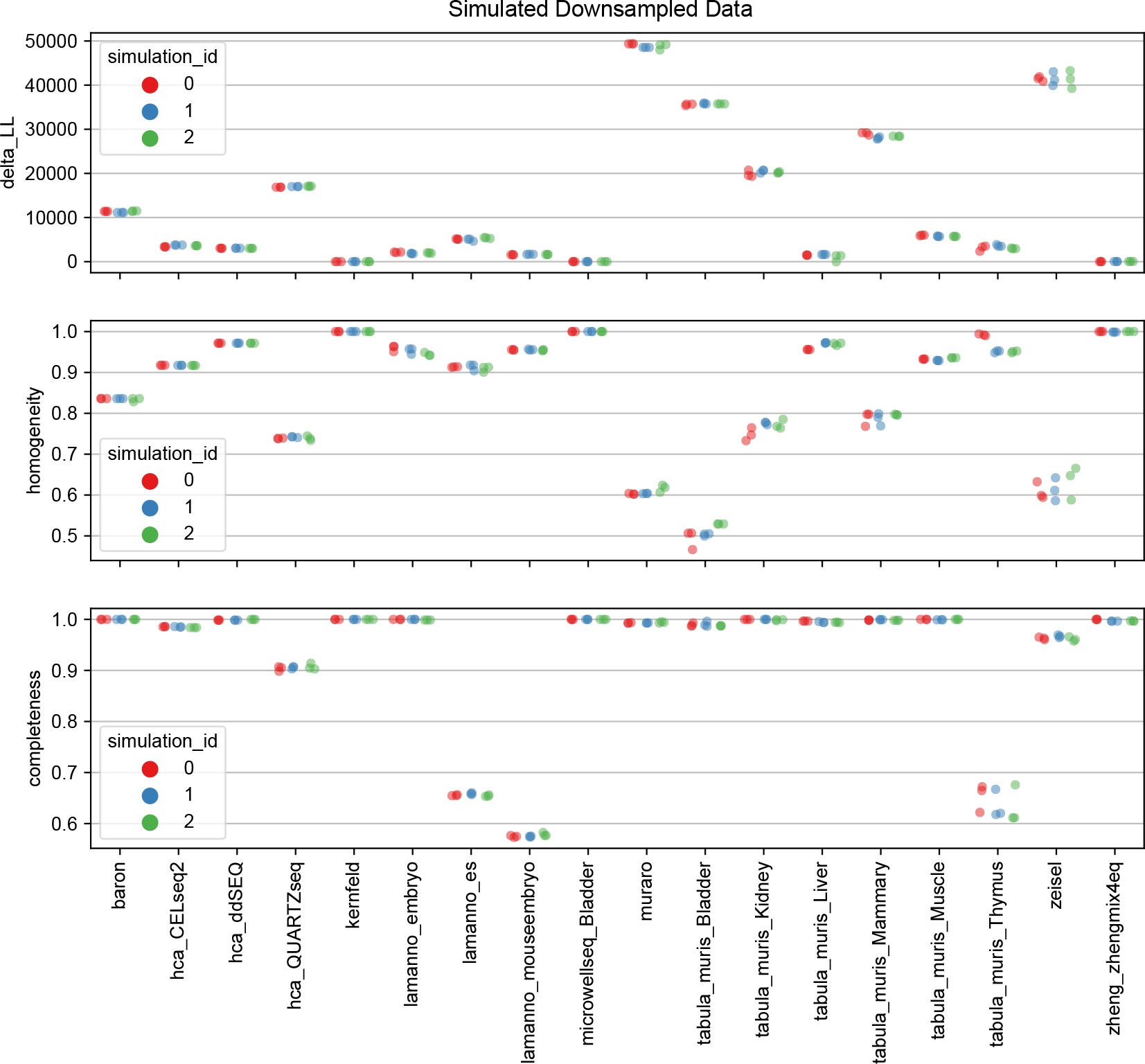
Detailed results from Cellstates runs on down-sampled simulated data based on the set of GESs from various indicated datasets, but with a median of only 1000 UMI per cell. For each of the 18 GES-sets, three simulations were generated (red, blue, and green) and Cellstates was run three times on each simulation. In the top panel, the difference in log-likelihood delta_LL between the inferred partition and the partition used to generate the simulated data is shown (positive meaning that a partition was found with higher log-likelihood than the one used to generate the data). The corresponding homogeneity and completeness of the inferred partitions compared to the partitions used to generate the data are shown in the two lower two panels.

**Figure S3:**
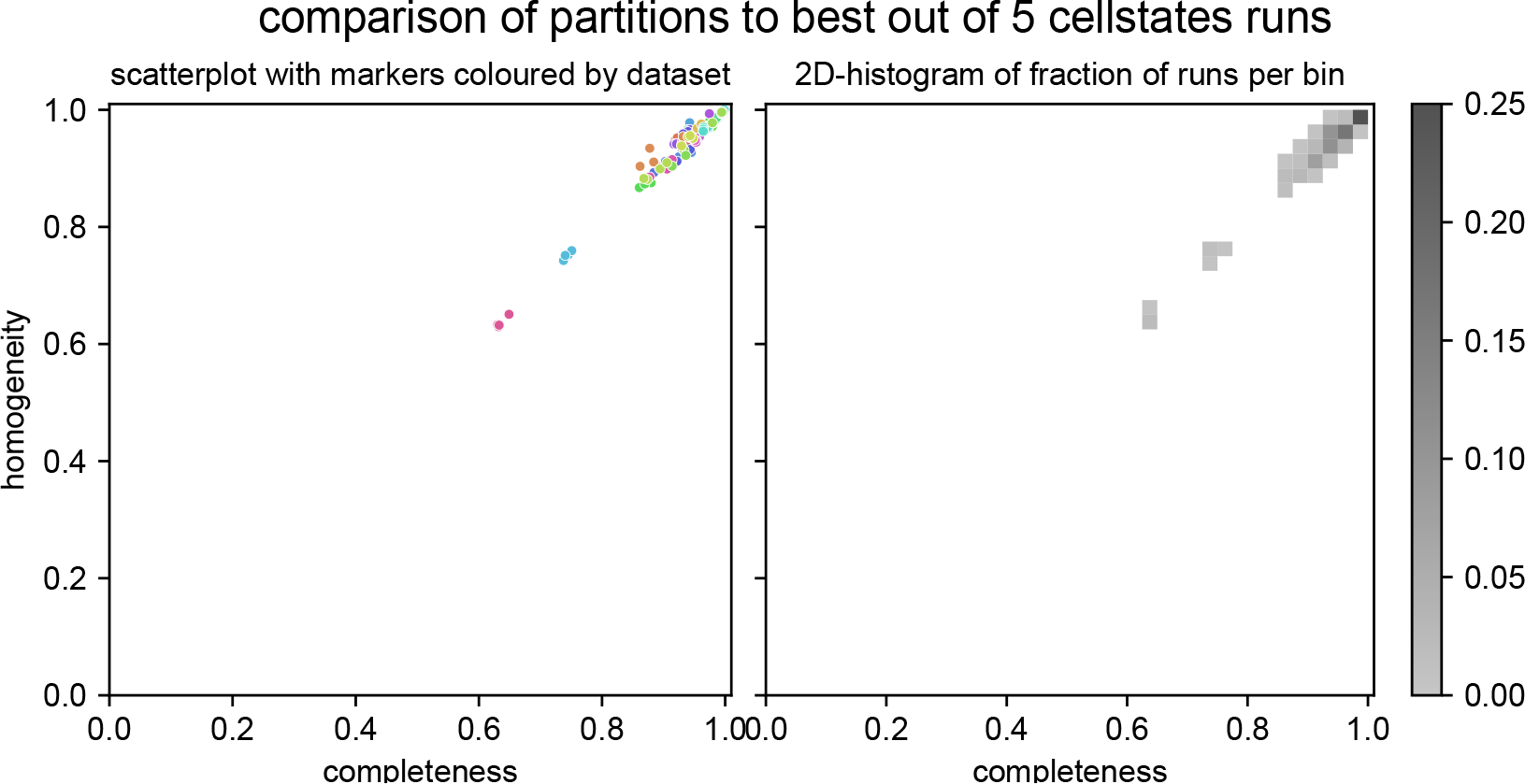
Reproducibility of independent Cellstates runs. Cellstates was run 5 times on each of 34 scRNA-seq datasets and the highest-likelihood partition that was found was compared with those of the 4 other runs. The resulting homogeneity and completeness scores are shown twice. On the left as a scatter-plot with markers colored by dataset, illustrating which outliers belong to the same dataset. On the right, a 2D-histogram shows the fraction of runs in bins of size 0.025 × 0.025, showing that most fall within the top right corner with homogeneity and completeness > 0.95.

**Figure S4:**
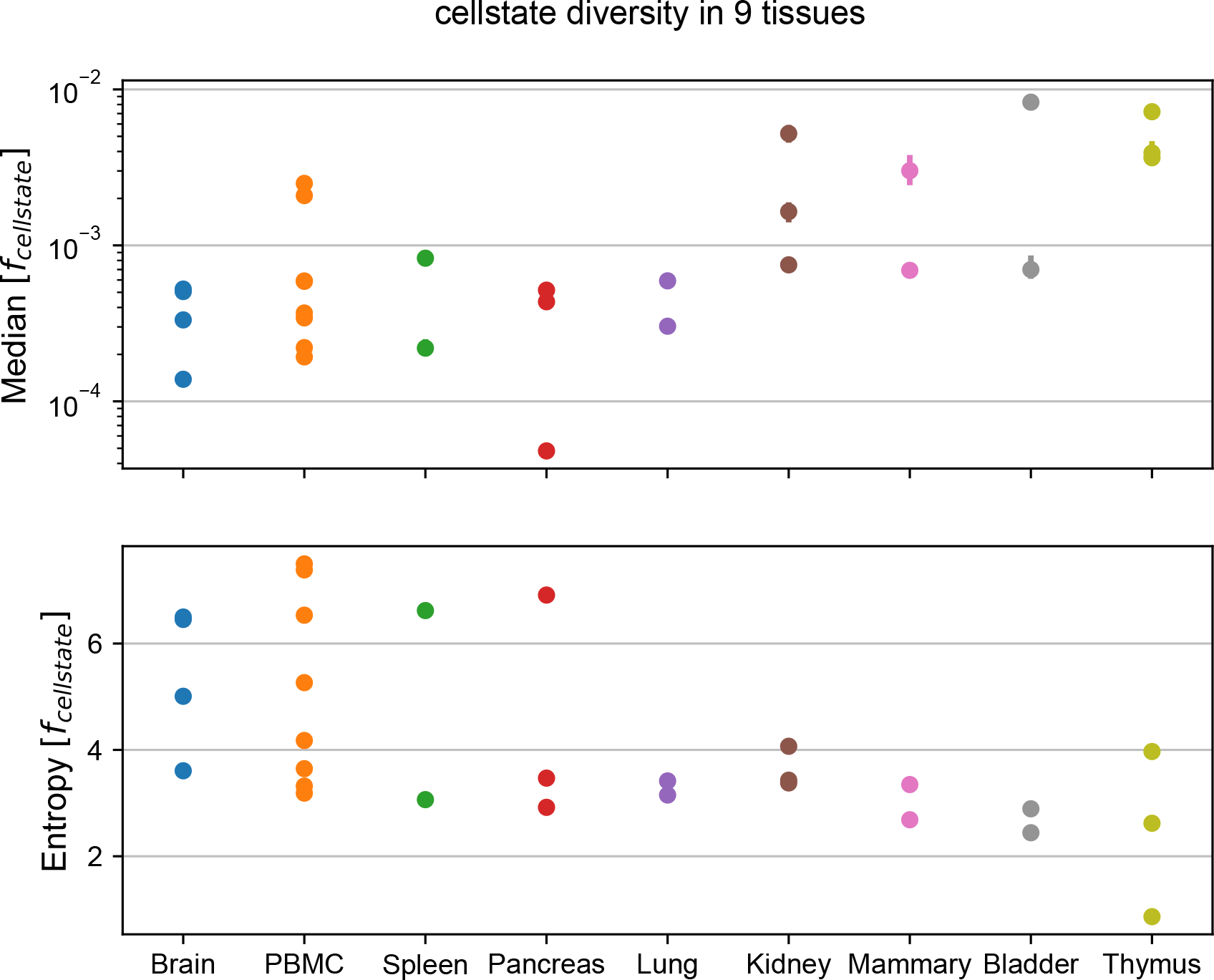
Tissue of origin is predictive for cellstate diversity. For 29 different datasets from 9 tissues the entropy of the distribution of cellstate abundances *f*_cellstate_ in each dataset are shown by tissue. Although there is large variability across datasets, datasets from the same tissue tend to have similar cellstate diversity. Tissues are sorted roughly from most to least diverse from left to right.

**Figure S5:**
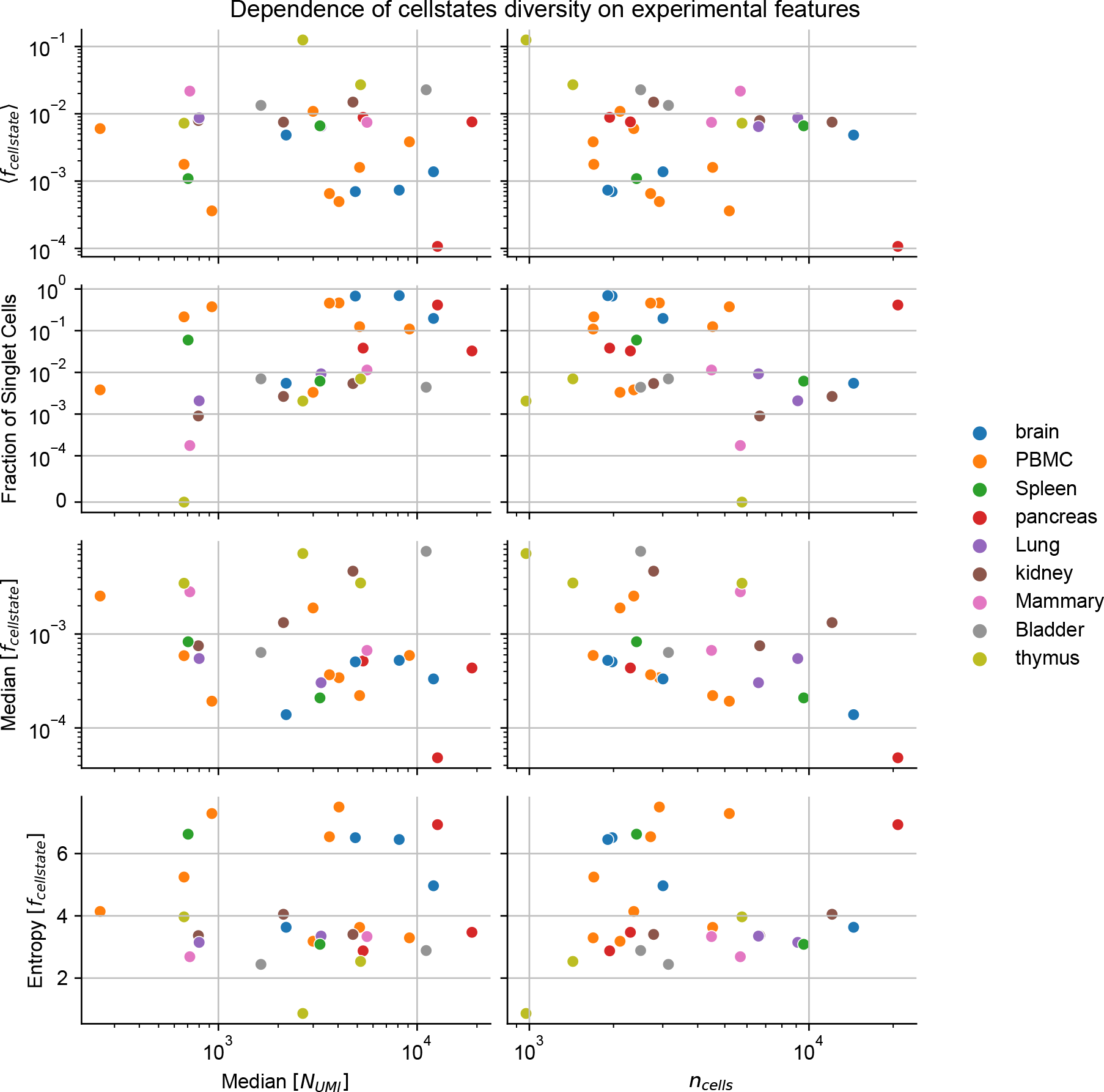
Cellstate diversity does not depend on technical features of the experiment. The mean of the distribution of cellstate abundances *f*_cellstate_ (top row panels), the fraction of singlet cellstates (second row panels), the median (third row panels) and the entropy (bottom row panels) of *f*_cellstate_ are shown as a function of the median total number of UMIs per cell (*N*_*UMI*_) (left column panels) and the total number of cells in each dataset (right column panels). Marker colors indicate the tissue from which the samples originate. These results show that there is no correlation between the various diversity measures and either sequencing depth (total UMI count) or the number of cells sequenced, with the exception of a weak negative correlation between the median cellstate abundance Median[*f*_cellstates_] and the number of cells *n*_cells_. This weak correlation is explained by the fact that the majority of cellstates are singlets in many experiments. Note that the variation in diversity measures for data from the same tissue (i.e. dots of the same color) along the *y*-axes is much less than that along the *x*-axes, which means that cellstate diversity is largely driven by biological and not technical experimental features.

**Figure S6:**
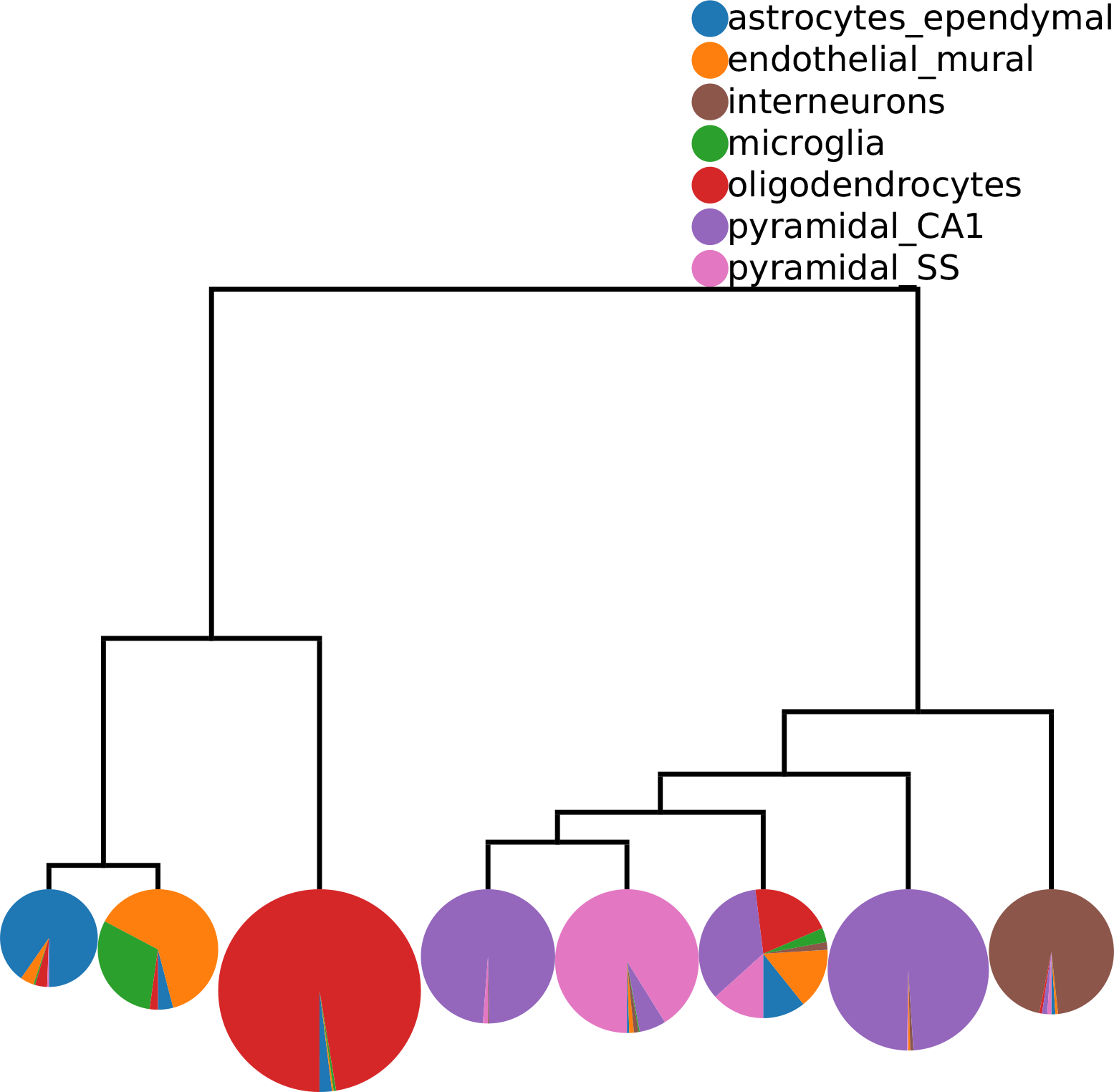
Cells in the 8 major cellstate clusters of the Zeisel dataset [19] largely match annotations provided in that publication. Colors indicate different annotations from [19] (see legend), each pie chart corresponds to one higher-order cluster, and the area in each pie chart is proportional to the number of cells with the corresponding annotation in that cluster. Note that cells annotated by [19] as microglia and endothelial-mural cells are merged into one cluster at this level of the hierarchy. The tree structure indicates how these clusters are related upon further merging.

**Figure S7:**
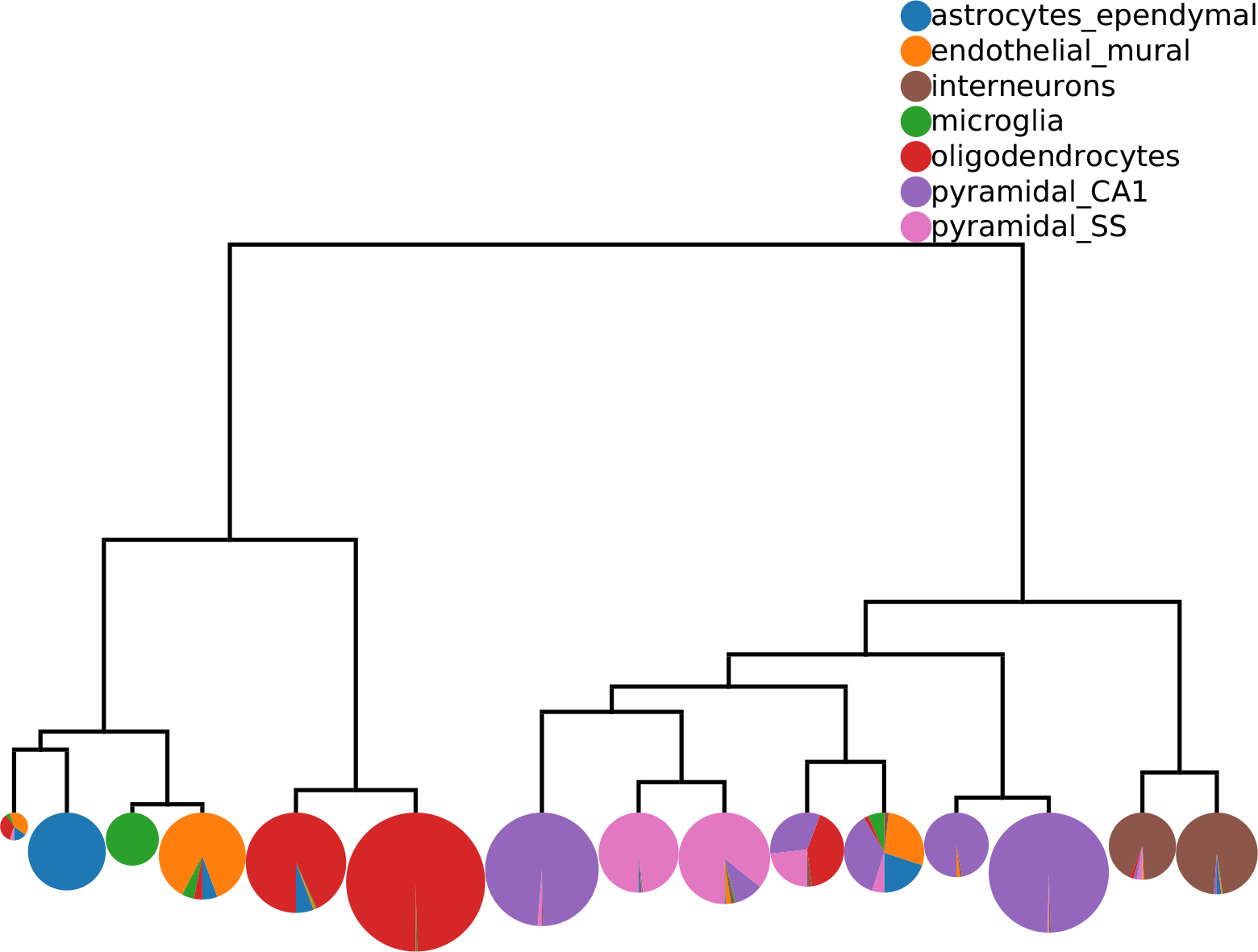
At 15 higher-order cellstate clusters cells annotated in [19] as microglia and endothelial-mural separate. Colors indicate different annotations from [19] (see legend) and the area in each pie chart is proportional to the number of cells in the corresponding cluster with the corresponding annotation. Note that at this level of the hierarchy cells with a common annotation in [19] tend to separate into multiple higher-order cellstate clusters.

**Figure S8:**
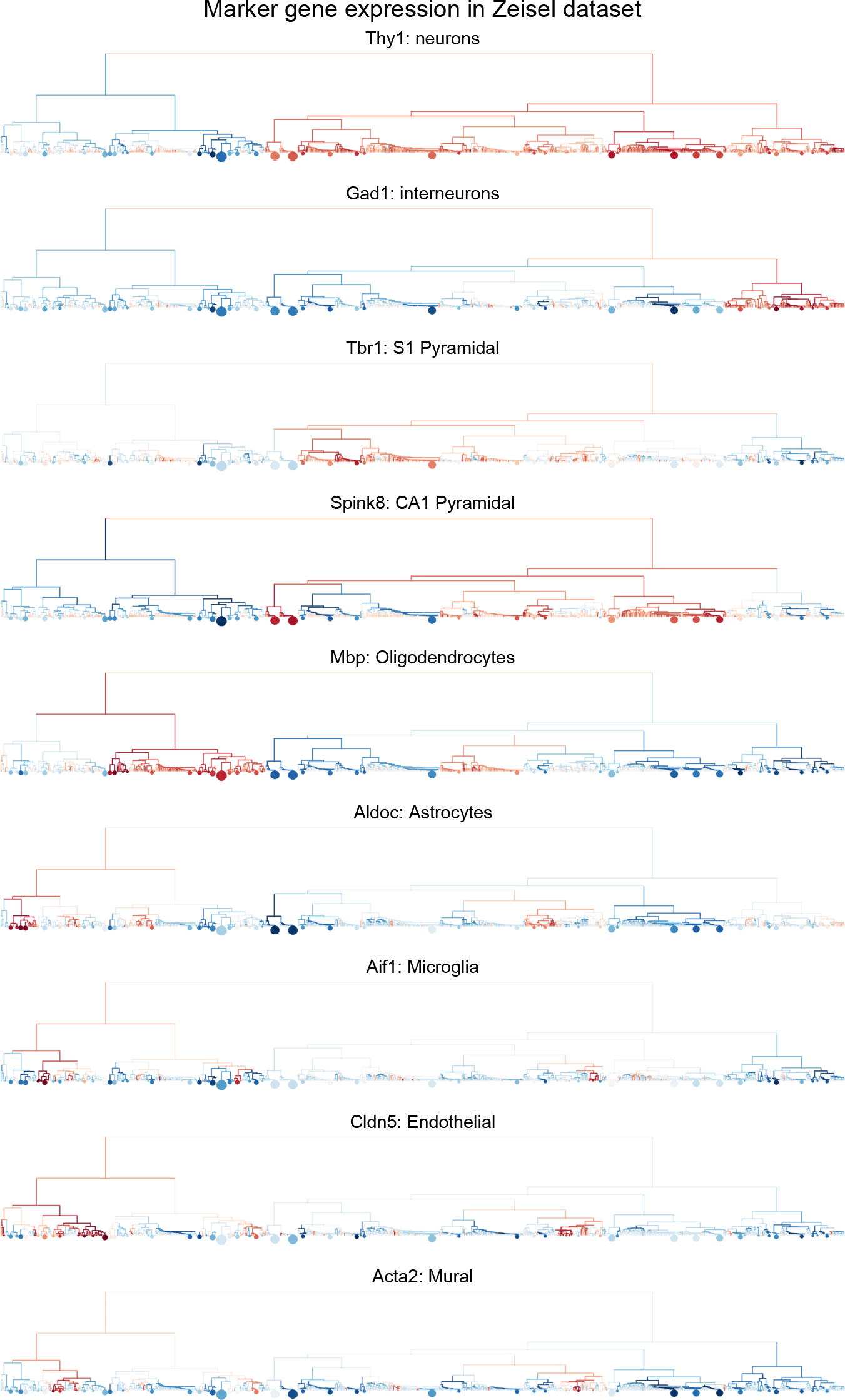
Expression of marker genes for annotated cell types visualized in the hierarchical tree of higher-order cellstate clusters. The sizes of the discs at the leaves correspond to the numbers of cells in the corresponding cellstates. See Figure 4 to compare with the higher-order clusters of Cellstates.

**Figure S9:**
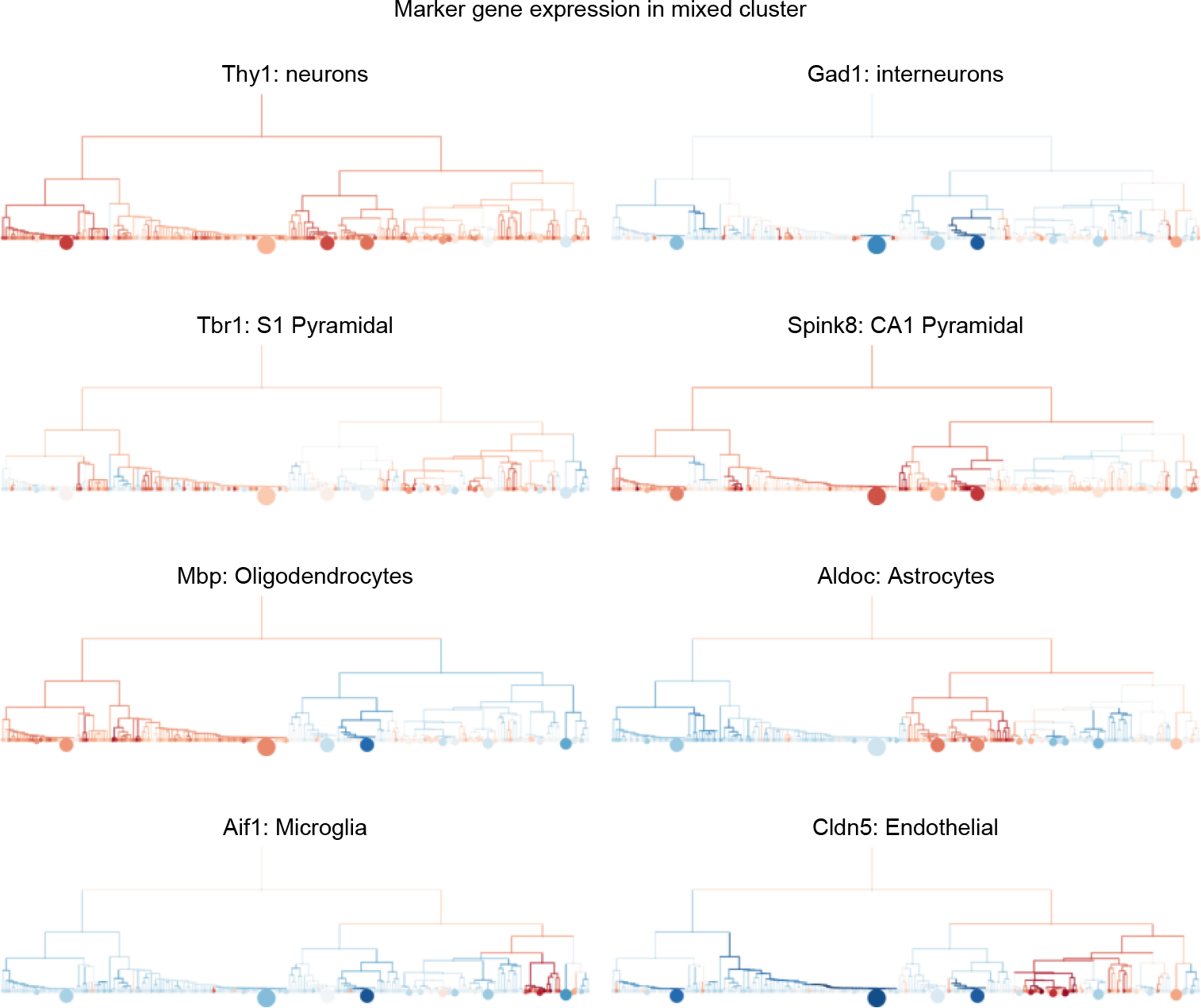
Expression of marker genes for annotated cell types visualized in the hierarchical tree of higher-order cellstate clusters. The sizes of the discs at the leaves correspond to the numbers of cells in the corresponding cellstates. See Figure 4 to compare with the higher-order clusters of Cellstates.

